# 5′-triphosphate guanosine RNAs recruit GTP-binding proteins to suppress RIG-I/IFN type I signaling

**DOI:** 10.1101/2023.12.22.573000

**Authors:** Jacek Szymanski, Magdalena Wolczyk, Ivan Trus, Zara Naz, Nathalie Idlin, Justyna Jackiewicz, Elzbieta Nowak, Christine Wuebben, Gunther Hartmann, Juri Rappsilber, Gracjan Michlewski

## Abstract

The interferon (IFN) response is crucial for antiviral activity, but its overstimulation can lead to a wide range of autoimmune disorders. The cytoplasmic pattern recognition receptor RIG-I detects viral double-stranded RNAs (dsRNAs) and endogenous polymerase III transcripts carrying a 5′-triphosphate (5′-ppp) or 5′-diphosphate (5′-pp) moiety, triggering phosphorylation of IRF3 and IFN immune response. While many viral RNAs initiate with 5′-ppp-adenosine (5′-pppA) and most endogenous Pol III transcripts in higher eukaryotes start with 5′-ppp-guanosine (5′-pppG), no apparent reason for this bias has been identified so far. Here we demonstrate that dsRNAs initiating with 5′-pppA trigger stronger RIG-I/IFN response than those starting with 5′-pppG. We show that several GTP-binding proteins interact preferentially with 5′-pppG RNAs. Finally, supplementation with guanosine, but not adenosine, which rapidly increases intracellular concentrations of GTP and ATP, respectively, eliminates the difference in immunogenicity between 5′-pppG and 5′-pppA RNAs. Our findings suggest that 5′-pppG RNAs may enable certain RNA viruses and Pol III transcripts to limit detection by innate immune receptors. These results offer new insights into the sequence-dependent activation of the RIG-I/IFN pathway and have important implications for both antiviral immunity and the role of Pol III-derived RNAs in autoimmune diseases.

## Introduction

In higher eukaryotes, the innate immune system acts as a rapid and essential defence mechanism against viral infections. This response is initiated when cellular sensors, known as Pattern Recognition Receptors (PRRs), detect molecular signatures commonly associated with pathogens, referred to as Pathogen-Associated Molecular Patterns (PAMPs). ^1^ One such molecular signature is the 5′-triphosphate (5′-ppp) group, which marks uncapped RNAs typically produced by viruses, but also present in some transcripts generated by RNA polymerase III. ^2,3^ Among the PRRs, Retinoic Acid-Inducible Gene I (RIG-I) plays a central role in detecting viral RNA that carries 5′-ppp motifs. Binding of such RNA induces a structural rearrangement in RIG-I and promotes its K63-linked ubiquitination by the E3 ligase Riplet. ^4,5^ This modification initiates a signaling cascade via the mitochondrial antiviral signaling protein (MAVS), leading to the activation of transcription factors IRF3, IRF7, and NF-κB. ^6^ Once activated, these factors migrate into the nucleus and drive the production of type I interferons, which initiate a broad antiviral program by upregulating interferon-stimulated genes (ISGs). ^7^ While essential for antiviral defence, uncontrolled activation of this pathway can result in chronic inflammation or autoimmunity. ^8^

RIG-I protein displays highly selective ligand recognition, responding to both structural and chemical features of RNA. Its activation is triggered by short double-stranded RNAs (dsRNAs) with blunt ends, as well as by distinct molecular signatures such as 5′-triphosphate (5′-ppp) or 5′-diphosphate (5′-pp) groups. ^9–12^ Crucially, RIG-I can distinguish these ligands from host RNAs by recognizing the absence of 2′-O-methylation, a modification typically found in the capped ends of eukaryotic mRNAs. ^13^ Even minimal synthetic RNA constructs, including 10-base-pair duplexes or stem-loop structures, are sufficient to robustly activate RIG-I. ^14,15^ During Influenza A virus infection, short viral RNA fragments approximately 80 nucleotides in length have been identified as highly potent RIG-I agonists, underscoring the critical role of RNA structure and composition in the activation of immune response. ^16^

It has been noted that a substantial number of RNA viruses’ genomes begin with 5′-ppp adenosine (5′-pppA) ^17^ and Pol III transcripts of higher eukaryotes and the genomes of some highly pathogenic RNA viruses initiate with 5′-ppp guanosine (5′-pppG). ^18–22^ To revisit this notion, we collated 5′-terminal nucleotides of representative RNA viruses, which are known to have 5′-ppp (Figure 1A and 1B). This collection revealed that some RNA viruses (such as influenza A virus - IAV) start predominantly from 5′-pppA, whereas other pathogenic viruses (including Ebola, Lassa, and HCV) can start from 5′-pppG or other nucleosides. Notably, attempts of mutations of the first nucleotide in viral genomes have already been noted in the literature. Intriguingly, IAV reverts to its wild-type configuration upon mutation or deletion of the first nucleotide, ^23^ while modifying the 5′ terminal nucleotide to another purine nucleotide in hepatitis C virus (HCV) is not feasible. ^21,24^ Moreover, HCV genotypes starting with 5′-pppA are largely capped with flavin adenine dinucleotide (FAD), which might prevent them from inducing RIG-I, while genotypes starting with 5′-pppG remain uncapped. ^21^

**Fig. 1.**
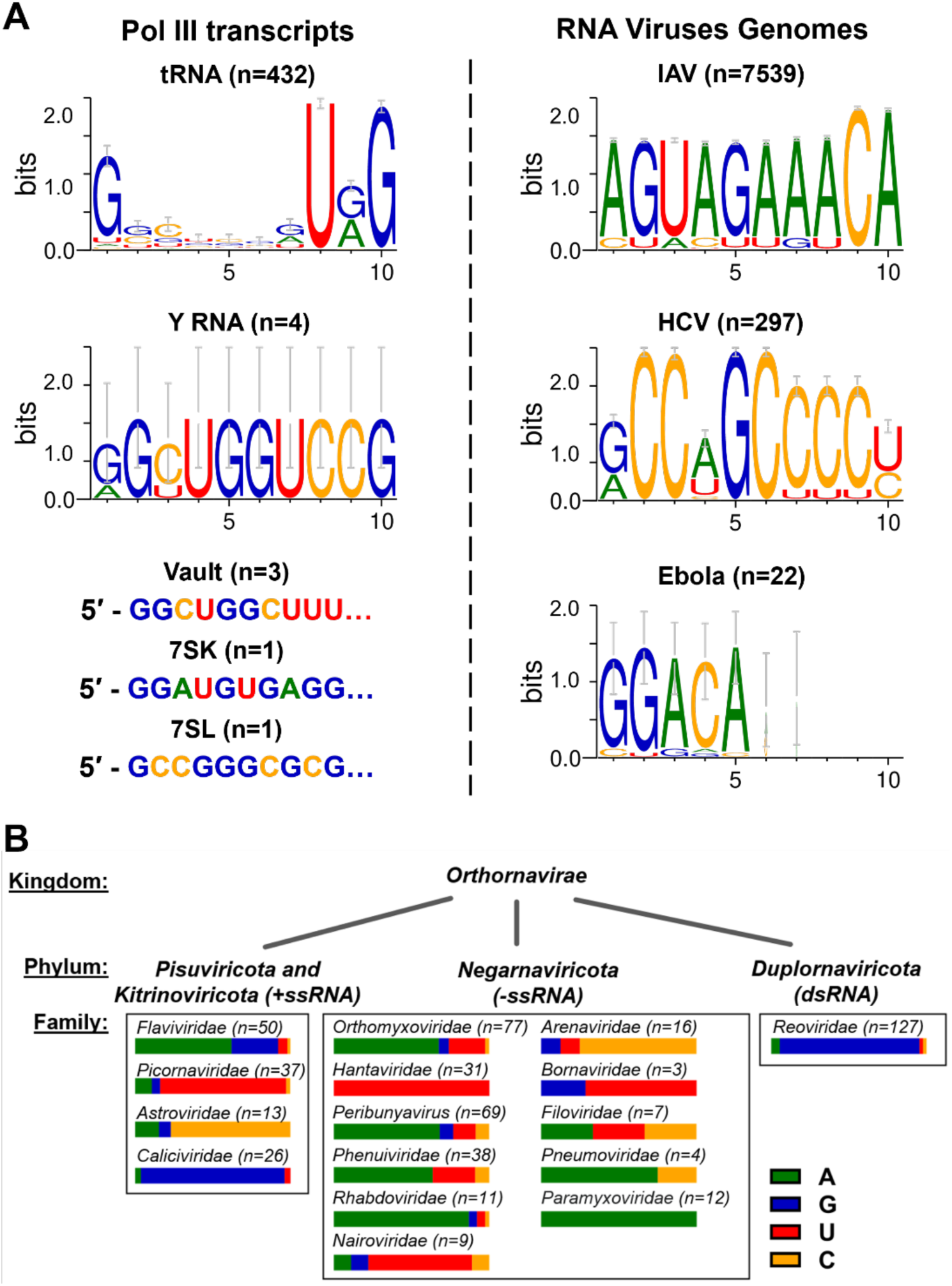
Terminal 5′-ppp nucleotide conservation in viruses and Pol III transcripts. (**A**) The conservation of the terminal 5′-ppp nucleotide in both Pol III transcripts and selected human viruses with uncapped RNA genomes. Nucleotide sequences representing the 5′ end of specific Pol III transcripts and selected RNA viruses. The sequence probability logos were generated using WebLogo 3. ^84^ The data sources include GenBank, tRNAscan-SE Genomic tRNA Database (http://gtrnadb.ucsc.edu/), and for Ebola sequences reported by Deflubé *et al.*_22_ (**B**) Variability in the 5′ nucleotide of human viruses in virus families with uncapped RNA genomes. The reference sequences of viral genomes were downloaded from the NCBI Viral Genome Browser (https://www.ncbi.nlm.nih.gov/genome/viruses/). The numbers in parentheses indicate the number of genomes/segments represented per family. The frequencies of initial nucleotides are depicted using horizontal bars.

On the other hand, Pol III transcripts are known to be synthesized with 5′-ppp (Figure 1A). Interestingly, most Pol III transcripts in higher eukaryotes, including non-coding tRNA, Y RNA, vault RNA, and 5S RNA, initiate from 5′-pppG. Although those transcripts are suboptimal RIG-I agonists with their 3′ end uridine-rich tails, they have been shown to be endogenous triggers of the RIG-I/IFN pathway. ^25,26^ Notably, among four human Y RNAs, only Y5 RNA starts from 5′-pppA, while the three others start from 5′-pppG. It has been reported that Y5 RNA shows mainly nuclear localization, ^27^ and our qRT-PCR (quantitative reverse transcription and polymerase chain reaction assay)-based quantification shows that Y5 RNA levels are the lowest compared to other Y RNAs in HEK293 cells (Supplementary Figure S1). The reasons for these sequence preferences in both viral RNA genomes and Pol III transcripts remain unknown.

Our goal was to investigate the impact of the first nucleotide on RIG-I pathway activation. Since viral RNA genomes cannot be readily mutated at the 5′ end, the most suitable alternative was to use synthetic RNAs, which can be easily manipulated. Currently, the most widely used method for RNA production in both research and the RNA-based drug industry is T7 polymerase-based *in vitro* transcription (IVT). While this technique offers several advantages, such as relatively low cost and high production efficiency, it is also associated with the generation of various side products. ^28^ These include transcripts resulting from random priming of abortive products, read-through transcription of run-off transcripts, and promoter-independent transcription from either the sense strand of RNA or the antisense strand of the DNA template. ^29–33^ Although numerous purification strategies have been proposed to eliminate IVT-derived contaminants (reviewed in ^34^), even RNAs purified by urea-PAGE will still contain highly immunogenic impurities formed by perfectly complementary RNA duplexes and the main RNA product. Notably, we have shown that IVT-based production of 5′-pppA generates much more immunogenic dsRNA than 5′-pppG. ^35^ Altogether, this makes the IVT-based preparation of single-stranded RNAs unsuitable for testing differences in immunogenicity between 5′-pppA and 5′-pppG transcripts.

To address this issue, we developed a strategy that utilizes only perfect dsRNA duplexes, produced using two different methodologies depending on the transcript length. For RNA molecules shorter than 40 nucleotides, we employed chemical solid-phase synthesis in two batches, in which the first strand, called “sense,” was synthetized with a 5′-ppp moiety and the second (“antisense”) lacked any phosphate modification (5′-OH). For transcripts longer than 40 nucleotides, we used IVT reactions to prepare both sense and antisense strands separately with each strand containing a 5′-ppp moiety. Our approach of using perfectly complementary dsRNA duplexes has two major advantages: first, the dsRNA products are highly potent RIG-I agonists, and second, preparation of IVT-derived dsRNA by mixing sense and antisense strands in 1:1 ratio eliminates the confounding effects caused by a tiny fraction of dsRNA impurities overrepresented in the 5′-pppA RNAs. ^35^

Here we provide the evidence that dsRNAs commencing with 5′-pppA elicit a considerably more robust RIG-I/IFN response in comparison to those originating with 5′-pppG. Notably, both dsRNAs exhibit comparable binding affinity and similar RIG-I activation in isolated systems, suggesting that differences in stimulation of the RIG-I/IFN pathway are unlikely to result from variations in their direct interaction with the receptor. Using RNA pull-down combined with mass spectrometry, we demonstrate that several GTPases and GTP-binding proteins exhibit a specific binding affinity for 5′-pppG RNAs. Finally, supplementation with guanosine which promptly elevates intracellular GTP levels through the nucleoside salvage pathway, but not with adenosine which increases ATP levels, abolishes the difference between 5′-pppG and 5′-pppA RNAs immunogenicity. We hypothesize that the highly abundant GTPases and GTP-binding proteins cause steric hindrance, thereby reducing RIG-I-mediated activation of the IFN response induced by 5′-pppG RNAs. Our findings suggest that 5′-pppG RNAs may enable certain RNA viruses and Pol III transcripts to evade detection by cellular immune sensors, uncovering a novel mechanism controlling sequence-dependent RIG-I/IFN pathway activation.

## Results

### 5′-pppA dsRNAs stimulate the RIG-I/IFN pathway stronger than 5′-pppG dsRNAs in human and murine cells

To evaluate the biological significance of the observed 5′ terminal nucleoside identity in RIG-I agonists, we hypothesized that retention of the 5′-pppG RNAs may be an adaptation that allows some RNA viruses and Pol III transcripts to evade detection by the RIG-I/IFN pathway. As stated in the introduction, direct mutagenesis of viruses or the use of IVT reactions to generate cognate single-stranded Pol III transcripts with similar levels of immunogenic dsRNA was not achievable. Instead, we employed two IAV-derived dsRNAs, which sense strand represents the beginning of the positive strand of segment 8^th^ of the IAV PR8 strain – 76 bp IVT-produced dsRNA (Figure 2A) and its shorter analogue 24 bp dsRNA produced by chemical synthesis (Figure 2B). To investigate the role of the terminal nucleoside, we replaced the initial 5′-pppA with 5′-pppG in both IVT produced and synthetic dsRNAs, while maintaining the base pairing between the 5′-end of the first strand and 3′-end of the second strand by creating Watson-Crick base pairs (A:U and G:C) for the initial 5′-pppA or 5′-pppG (Figures 2A and 2B). The quality of dsRNA production was assessed through a two-step evaluation process. First, the integrity of the individual RNA strands was analyzed using denaturing urea-polyacrylamide gel electrophoresis (urea-PAGE) (Supplementary Figures S2A and S2C). Subsequently, successful duplex formation was verified via native polyacrylamide gel electrophoresis (native PAGE) (Supplementary Figures S2B and S2D). Then, HEK293 human embryonic kidney cells were transfected with either IVT 76 bp and synthetic 24 bp dsRNAs bearing 5′-pppA or 5′-pppG as a 5′ terminal nucleotide. The IVT 76 bp dsRNAs were tested at concentrations ranging from 0.1 ng/ml to 100 ng/ml, while the synthetic 24 bp dsRNAs were tested between 1 ng/ml and 1000 ng/ml. After 24 hours post transfection, IRF3 phosphorylation and RIG-I protein levels were assessed by western blot analysis. In cells treated with IVT 76 bp dsRNAs, higher levels of phosphorylated IRF3 (pIRF3) and RIG-I protein were observed in 5′-pppA dsRNA-transfected cells compared to 5′-pppG dsRNA-transfected cells, particularly at concentrations between 0.1 and 10 ng/mL (Figure 2C). The difference in pIRF3 and RIG-I was observed in all tested time points from 4 up to 24 hours post transfection (Supplementary Figure S3A). Moreover, the effects of varying strength of RIG-I signaling activation were also observed at the mRNA level of *IFN1β* and *ISG15* (Interferon-Stimulated Gene 15) measured by qRT-PCR (Supplementary Figure S3B). A similar trend in RIG-I and pIRF3 expression was observed in cells transfected with synthetic 24 bp dsRNAs, although the effect was shifted to higher concentrations. Enhanced pIRF3 and RIG-I levels for 5′-pppA over 5′-pppG 24bp dsRNA were detected for 10 and 100 ng/ml concentrations, whereas no signs of RIG-I stimulation were observed at 1 ng/ml (Figure 2D). IVT-derived dsRNAs appeared to be more stimulatory for the RIG-I pathway, which may be attributed to a higher length but also the presence of 5′-ppp moieties on both ends of the RNA duplex in contrast to synthetic dsRNAs, which carry a 5′-triphosphate only on one strand. At higher dsRNA concentrations (10 ng/ml for IVT dsRNA and 100 ng/ml for synthetic dsRNA), a saturation effect was observed, with no observed difference in pIRF3 and RIG-I levels between 5′-pppA and 5′-pppG (Figures 2C and 2D).

**Fig. 2.**
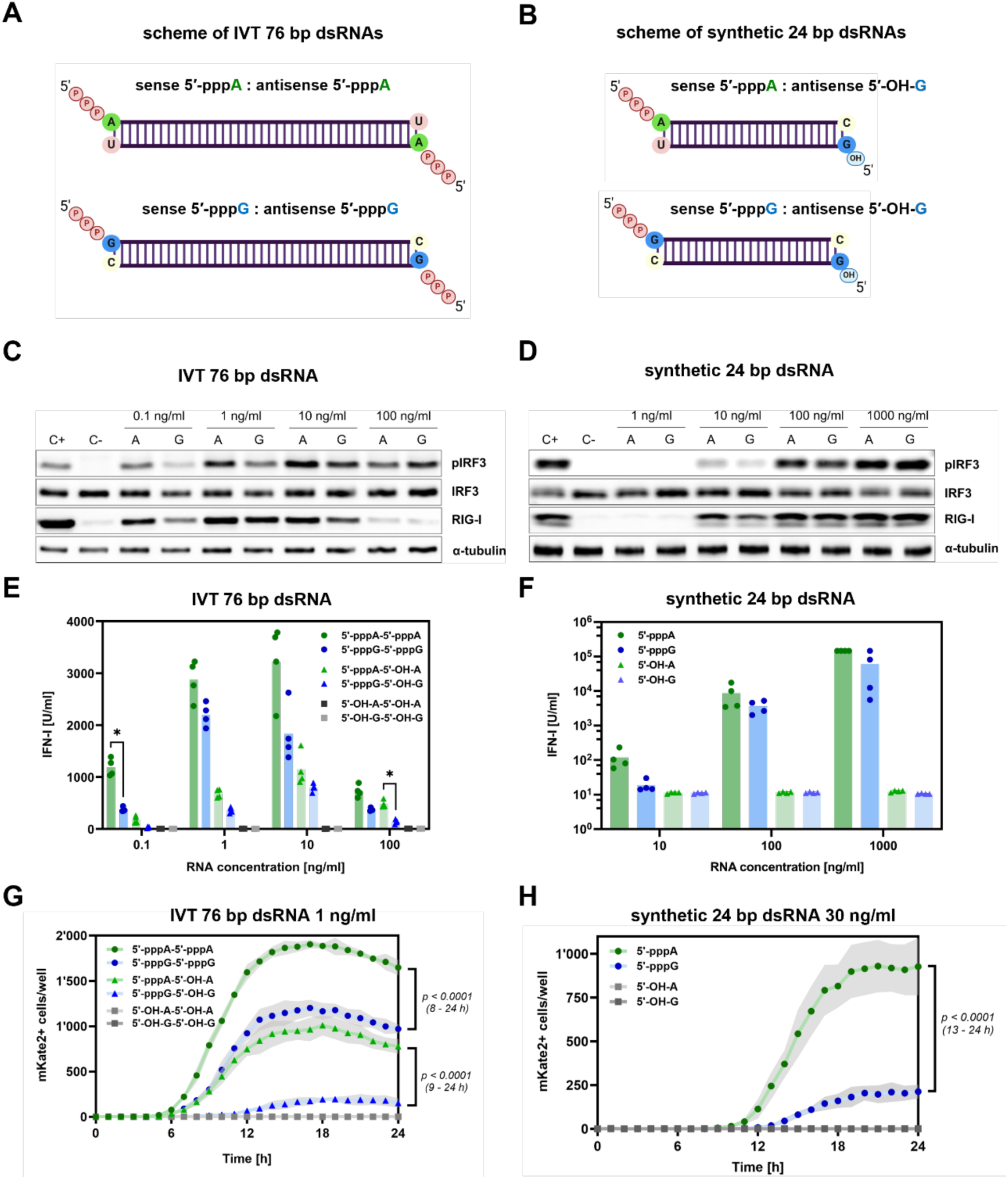
5′-pppA dsRNA promotes more robust RIG-I/IFN signaling than 5′-pppG. (**A, B**) Schematic representation of dsRNA variants differing at the 5′ terminal nucleotide (5′-pppA or 5′-pppG) produced via in vitro transcription and chemical synthesis, respectively. (**C, D**) Western blot analysis of IRF3 phosphorylation and RIG-I protein levels in HEK293 cells treated with varying concentrations of 76 bp IVT (**C**) or 24 bp synthetic dsRNAs (**D**). (**E, F**) Type I interferon production in HEK293 cells was assessed with HEK-Blue IFN type I assay after transfection with dsRNAs. Two-way ANOVA with Šídák’s multiple comparisons test was used for statistical analysis. (**G, H**) RIG-I/IFN activation in murine BMDMs upon transfection with tested dsRNAs, shown as mKate2+ cell counts over 0–24 h post-transfection. Grey area represents standard deviation for n=5. Data were analyzed using repeated measures two-way ANOVA on log-transformed values [log_10_(x + 1)], with Geisser-Greenhouse correction and Šídák’s multiple comparisons test.

Using IFN-α/β reporter HEK293 cells and colorimetric HEK-Blue IFN type I assay, we assessed the activity of IFN-α/β induced by the tested RNAs. Notably, IVT dsRNAs bearing a 5′-pppA induced significantly stronger IFN responses compared to those with 5′-pppG (Figure 2E). However, this difference reached statistical significance only at the lowest concentration tested (0.1 ng/ml). In case of synthetic dsRNAs, a trend of enhanced IFN production by the 5′-pppA variant was observed at 1 ng/ml, but no difference was observed in higher concentrations of dsRNA between the 5′-pppA and 5′-pppG variants (Figure 2F). Importantly, HEK293 cells transfected with either IVT 76 bp or synthetic 24 bp dsRNA lacking the 5′-triphosphate moieties showed no stimulation of IFN-α/β production (Figures 2E and 2F). Removal of 5′-triphosphate groups from the antisense strands of 76 bp dsRNA resulted in reduced phosphorylation of IRF3 and decreased expression of RIG-I for both 5′-pppA and 5′-pppG dsRNA variants (Supplementary Figure S3C). Notably, the differential response between the 5′-pppA and 5′-pppG forms remained evident. Complete dephosphorylation of both RNA strands in dsRNAs led to full suppression of the RIG-I/IFN signaling pathway (Supplementary Figure S3C).

To minimize potential false positive results due to IVT contaminants, through a semisynthetic approach we generated 76 bp dsRNAs. Specifically, a 55 nt RNA produced by IVT was ligated to a chemically synthesized 21 nt RNA using splint ligation to create a triphosphorylated sense strand, as previously described. ^35^ When evaluated in the HEK-Blue IFN type I assay, both 76 nt semisynthetic ligated 5′-pppA and 5′-pppG sense strand RNAs exhibited no detectable immunostimulatory activity at the tested concentrations (Supplementary Figure S3D). However, the addition of trace amounts (0.05–1%) of a fully complementary, IVT-produced, dephosphorylated 76 nt antisense RNA strand to form complete dsRNA was sufficient to activate the RIG-I/IFN pathway. Importantly, the distinct responses between 5′-pppA and 5′-pppG RNAs were still observable (Supplementary Figure S3D).

We next conducted transfection experiments using reporter bone marrow-derived macrophages (BMDMs) isolated from homozygous *mKate2* reporter mice, which enable real-time visualization of IFN-β expression via *mKate2* fluorescence. ^35^ A more robust IFN-β promoter activation was observed for RNAs initiating with 5′-pppA in cells transfected with either IVT-derived 76 bp dsRNA or fully synthetic 24 bp dsRNA (Figures 2G and 2H). In the case of 76 bp IVT dsRNA, a statistically significant difference in fluorescence between the 5′-pppA and 5′-pppG variants emerged at 8 hours post-transfection. For the synthetic 24 bp dsRNAs, this distinction became significant at 13 hours post-transfection. Throughout the time course, 5′-pppA RNAs were up to three times and fourteen times more immunogenic than the corresponding 5′-pppG constructs, for 76 bp and 24 bp dsRNAs respectively (Supplementary Figures S3E and S3F). Notably, no activation of the RIG-I/IFN signaling pathway was detected in BMDMs transfected with dsRNAs lacking 5′-triphosphate moieties, confirming the requirement of this structural feature for immune activation.

The observed effects were not limited to our two representative RNAs. Similar patterns were seen with IVT-derived dsRNAs, whose sense-strand sequences were taken from pre-let-7a and Y5 RNA (Supplementary Figures S4 and S5). Notably, the immunogenicity of Y5 RNA-derived dsRNA was strongly dependent on its concentration. At 0.1 ng/ml, the 5′-pppA Y5 dsRNA was up to forty times more immunogenic than the corresponding 5′-pppG construct, whereas at 10 ng/ml it was only about twice as immunogenic (Supplementary Figures S5F and S5G). Additionally, Y5-derived dsRNAs with 6xGC clamps at both ends exhibited substantially lower immunogenic potential than their wild type counterparts. Furthermore, a short synthetic, partially double-stranded RIG-I agonist with a blunt end (details available in the patent WO/2025/088117A1) also demonstrated that 5′-pppA RNA is up to twenty times more potent RIG-I stimulant than 5′-pppG RNA (Supplementary Figure S6). These results show that the increased immunogenicity of 5′-pppA-containing RNAs, compared with 5′-pppG, can be observed across a range of RIG-I agonists.

Taken together, these findings reveal a consistent pattern whereby dsRNAs bearing 5′-pppA ends are more immunogenic in both human and murine cells than equivalent sequences bearing a 5′-pppG terminus, highlighting the importance of RNA concentration in this context. Although this difference is observed in the human continuous cell line, a murine assay using primary cells (BMDMs with an *mKate2* reporter that allows real-time monitoring of IFN-β promoter activity) reveals a more robust difference in immunogenicity based on the identity of the 5′ terminal nucleotide. Furthermore, this effect is associated with the RIG-I/IFN signaling pathway, as stimulation is entirely reliant on the presence of 5′-triphosphate moieties.

### 5′-pppA and 5′-pppG RNAs show different binding affinity but similar activation in biochemical assays with recombinant RIG-I

To further investigate whether the observed significant difference in RIG-I/IFN pathway activation by 5′-pppA and 5′-pppG RNAs could be due to impaired binding of RIG-I to these RNAs, electrophoretic mobility shift analysis (EMSA) was performed. This assay was used to measure the binding affinities of 5′-pppA and 5′-pppG double-stranded RNAs to purified, recombinant RIG-I protein (Figures 3A and 3B and Supplementary Figure S7). Due to the method requirement of a fluorophore presence in the RNA molecule, 3′ end modification of the sense strand for IVT-produced dsRNA and 5′ end modification of the antisense strand with fluorescein amidite (6-FAM) for synthetic 24 bp dsRNA was introduced. Binding affinity measurements revealed that RIG-I binds to 5′-pppA IVT 76 bp dsRNA with significantly higher affinity (Kd = 1.47 nM) compared to 5′-pppG dsRNA (K_d_ = 2.32 nM) (Figure 3E and Supplementary Table S1). Notably, dephosphorylation of the 5′-pppA dsRNA resulted in an ∼8-fold reduction in RIG-I binding affinity (K_d_ = 10.84 nM), whereas the corresponding 5′-OH-G dsRNA exhibited only a modest, non-significant decrease in binding affinity (K_d_ = 2.94 nM) relative to its triphosphorylated counterpart. A similar trend was observed for direct interaction between recombinant RIG-I and synthetic 24 bp dsRNA (Figure 3B). The binding affinity for 5′-pppA (K_d_ = 0.40 nM) was significantly higher compared to 5′-pppG (K_d_ = 0.94 nM). Consistent with previous observations, the interaction of RIG-I with 5′-OH-A dsRNA was approximately 8-fold weaker (K_d_ = 4.50 nM) than with its triphosphorylated form and decrease in binding affinity to 5′-OH-G was reduced by only ∼2-fold (K_d_ = 1.81 nM) relative to 5′-pppG, but in this case the difference was significant (Figure 3E and Supplementary Table S1).

**Fig. 3.**
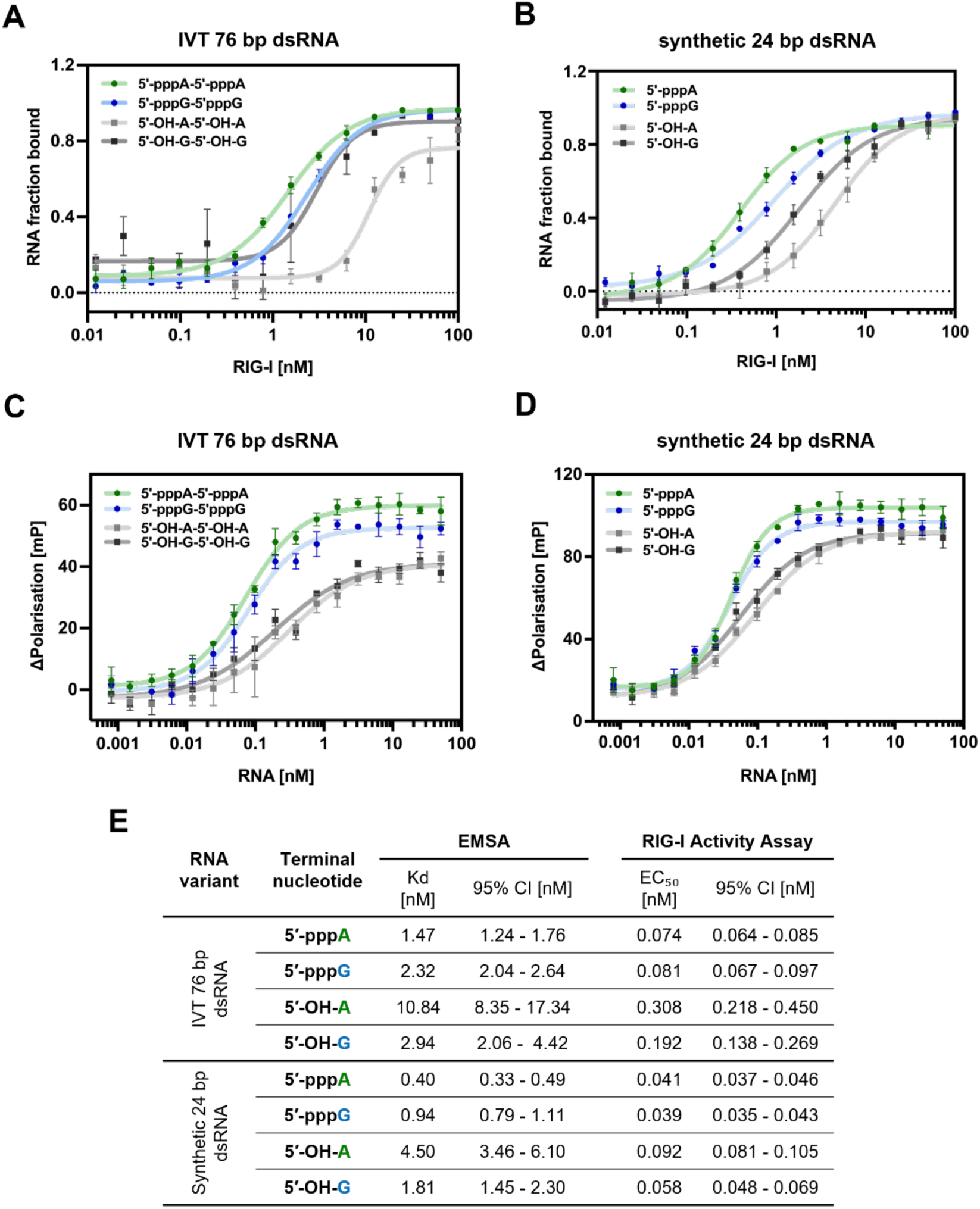
Biochemical analysis of recombinant RIG-I binding affinity and ATPase activity in response to dsRNA. (**A, B**) Binding affinity of recombinant RIG-I to dsRNAs with different 5′-terminal nucleotides (5′-pppA, 5′-pppG, 5′-OH-A, 5′-OH-G) was assessed using an electrophoretic mobility shift assay (EMSA). Plots represent the fraction of RNA bound to RIG-I at increasing protein concentrations. (**C, D**) ATP to ADP conversion was measured in the ATPase to assess recombinant RIG-I protein activation with dsRNAs. The 4PL model was fitted to EMSA and ATPase assay derived data to obtain dissociation constant (K_d_) for binding (**A, B**) and half maximal effective concentration of dsRNA (EC_50_) for ATPase activity (**C, D**). (**E**) Summary table of biochemical parameters, including K_d_ and EC_50_ values with corresponding 95% confidence intervals (CI), used to compare RIG-I binding affinity and activation across the different dsRNA variants.

To gain a deeper understanding of the interaction between RIG-I and dsRNA in relation to the 5′ terminal nucleotide under isolated conditions, we performed a RIG-I activation measurement. Based on previous studies that demonstrate a correlation between RIG-I activation by 5′-ppp RNA and its ability to hydrolyse ATP to ADP, an ATPase assay was applied to assess the activity of recombinant RIG-I upon dsRNA binding. A parameter used to compare RIG-I activity upon binding to different RNAs was the half maximal effective concentration (EC_50_) of the tested RNAs (Figures 3C and 3D). The result obtained for IVT 76 bp dsRNA by this methodology showed that triphosphorylated dsRNAs were most triggering RIG-I activity and differences between 5′-pppA (EC_50_ = 0.074 nM) and 5′-pppG (EC_50_ = 0.081 nM) were minimal and non-significant. Triphosphate-depleted dsRNAs were less effective ATPase activators, and in line with EMSA results, 5′-OH-A (EC_50_ = 0.308 nM) was much less RIG-I stimulatory than 5′-OH-G (EC_50_ = 0.192 nM). Similarly, synthetic 5′-ppp dsRNAs robustly activated RIG-I, with negligible differences between 5′-pppA (EC_50_ = 0.041 nM) and 5′-pppG (EC_50_ = 0.039 nM), and RNAs lacking triphosphate moiety – 5′-OH-A (EC_50_ = 0.092 nM) and 5′-OH-G (EC_50_ = 0.058 nM) – showed substantially reduced potency (Figure 3E and Supplementary Table S1).

Comparing those results with experiments performed on human and mouse cells, we speculate that immunogenic potential of dsRNA might be only partially explained by the difference in direct binding affinity of RIG-I protein to 5′-pppA and 5′-pppG dsRNA. However, the immunogenic potential of dsRNA does not strictly correlate with ability to activate RIG-I, as we observed no differences in ATPase assays. This suggests that the observed mechanism of differential immunogenic potential of 5′-pppA *vs.* 5′-pppG RNAs cannot be fully recapitulated in isolated conditions with recombinant RIG-I protein. Therefore, the difference in RIG-I pathway activation by 5′-pppA and 5′-pppG RNAs might be due to sequence-specific cellular factors that are not present in biochemical assays.

### 5′-pppA and 5′-pppG RNAs bind different sets of RBPs

To elucidate the cellular mechanism behind the difference in the immunogenicity of 5′-pppA *vs.* 5′-pppG RNAs, we employed RNA pull-down coupled with label-free quantitative mass spectrometry (RP-MS), ^36^ identifying proteins in cell extracts that exhibited differential binding affinities to the tested dsRNAs (Figure 4A). While most proteins displayed similar binding between the two RNAs, a subset of proteins emerged as outliers (Figure 4B). The 5′-pppA IVT 76 bp dsRNA pull-down enriched for a substantial group of proteins involved in nucleic acid metabolism, RNA stability, or degradation. Notably, the 5′-pppG IVT 76 bp dsRNA pulled-down proteins associated with translation and RNA transport. A considerable proportion of proteins enriched in the 5′-pppG IVT 76 bp dsRNA pull-down possesses GTP binding, small GTPase binding, or GTPase activity and are involved in translation process (Figure 4C). Conversely, few binders of 5′-pppA IVT 76 bp dsRNA were classified as single stranded RNA binding proteins, which is somewhat unexpected given that a dsRNA molecule was used as the bait in the pull-down assay (Figure 4D).

**Fig. 4.**
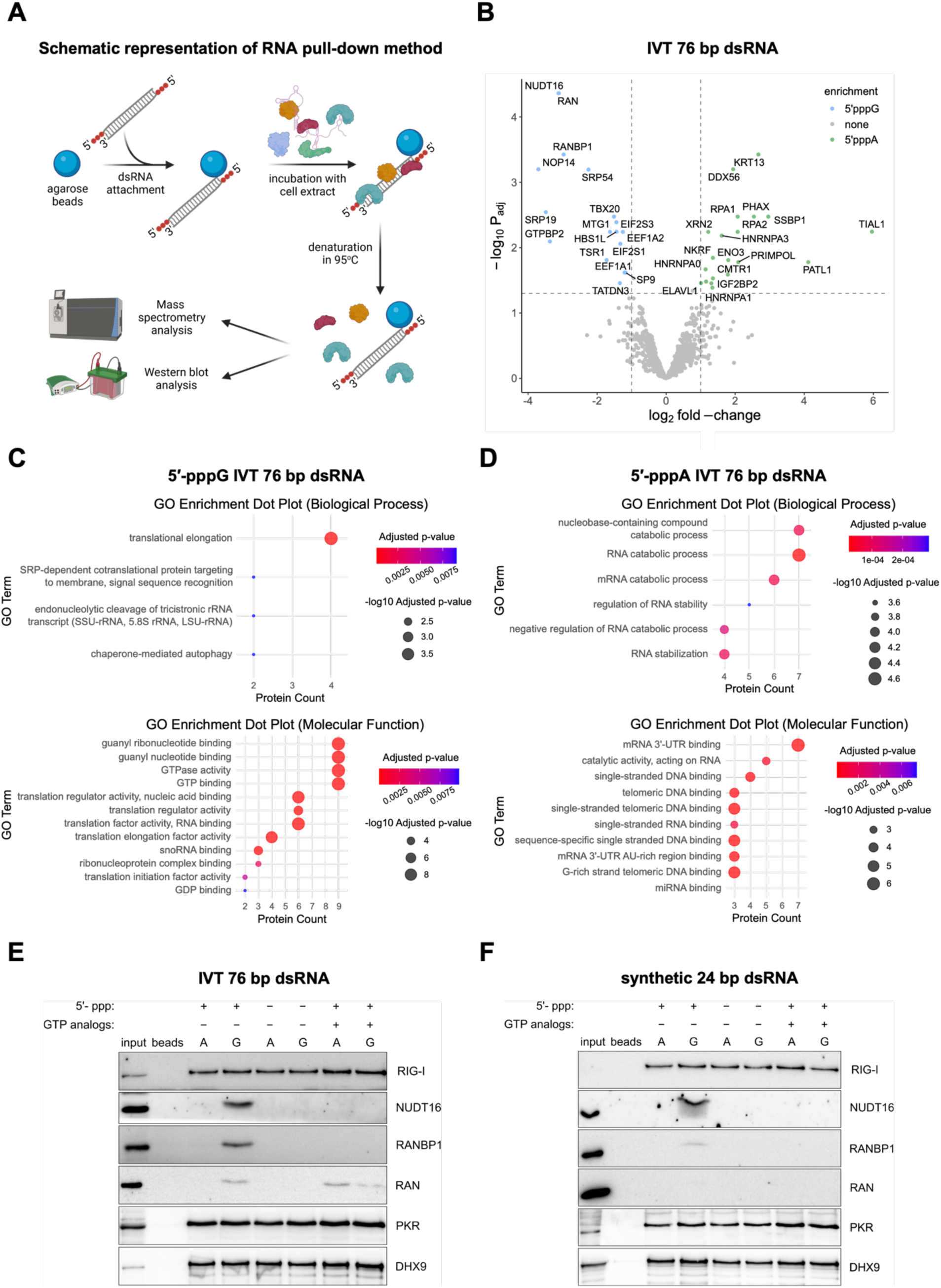
RNA pull-down analysis reveals RBP binding selectively to 5′-pppA or 5′-pppG dsRNA. (**A**) Schematic representation of RNA pull-down assay. (**B**) RNA pull-down followed by label-free LC-MS/MS was performed to investigate the interactome of IVT 76 bp dsRNA variants differing at the 5′ terminal nucleotide. Volcano plot represents differential analysis results of proteins enriched with dsRNA bearing 5′-pppA or 5′-pppG. Significance thresholds were set at |log_2_FC| > 1 and *p*_adj_ > 0.05. Proteins enriched with 5′-pppA are highlighted in green, while those enriched with 5′-pppG are shown in blue. (**C, D**) Gene Ontology (GO) analysis was conducted to identify the predominant biological processes and molecular functions associated with the differentially enriched proteins. (**E, F**) Binding of the top three candidate proteins – enriched on 5′-pppG IVT 76 bp dsRNA and associated with GTP-binding activity – was validated by RNA pull-down followed by Western blot analysis. Validation was performed using both IVT 76 bp (**E**) and synthetic 24 bp (**F**) dsRNA. PKR and DHX9 served as positive controls for dsRNA binding. Protein enrichment was further assessed under two conditions: depletion of the 5′-triphosphate moiety, and treatment with GTP nonhydrolyzable analogs, which acts as inhibitors of GTP-binding proteins.

To further validate these findings, we employed RNA pull-down western blot analysis using HEK293 cell lysates and either IVT 76 bp or synthetic 24 bp dsRNAs (Figure 4E and 4F). These results confirmed the increased binding of NUDT16, RAN, and RANBP1 proteins to the 5′-pppG IVT 76 bp dsRNA and its shorter analogue -synthetic 5′-pppG 24 bp dsRNA. Simultaneously, double-stranded RNA binding proteins PKR and DHX9 were captured by 5′-pppA and 5′-pppG RNAs in a similar way, confirming equal RNA quality and loading (Figures 4E and 4F). Surprisingly, no apparent difference in RIG-I binding between 5′-pppA and 5′-pppG RNAs was observed. This may indicate that RNA pull-down is not sensitive enough to detect subtle differences in protein affinity, or that the factors influencing differential RIG-I stimulation by 5′-pppA and 5′-pppG differ between *in cellulo* and *ex vivo* conditions. Finally, to check the dependence of binding of the identified proteins to the tested RNAs on the presence of the triphosphate moiety at the 5′ end of RNAs, we also tested dephosphorylated RNAs. Intriguingly, we observed that removing the 5′-ppp group from dsRNAs does not affect binding of RIG-I protein but results in complete inhibition of binding of 5′-pppG enriched proteins to dephosphorylated RNAs initiating with G (Figures 4E and 4F). Next, to investigate the influence of GTP binding proteins on the RIG-I-RNA interaction, we introduced non-hydrolyzable GTP analogs to protein extracts during the RNA pull-down procedure. This approach aimed to occupy the GTP-binding pockets, thereby impeding the interaction of the GTP-binding proteins with 5′-pppG RNAs. Indeed, supplementation with GTP analogs led to a complete blockade of NUDT16, RAN, and RANBP1 binding to RNAs (Figures 4E and 4F). In contrast, the binding of reference proteins DHX9 and PKR to dsRNAs remained unaffected. Taken together, these findings identify for the first time proteins that bind differentially to dsRNA containing 5′-pppG and 5′-pppA and show that GTP-binding proteins use a shared pocket to bind both GTP and 5′-pppG RNAs.

### Nucleoside salvage pathway-driven GTP increase equalises immunogenicity of 5′-pppG and 5′-pppA dsRNAs

To assess whether GTP-binding proteins have any effect on the activation of the RIG-I/IFN pathway *in cellulo*, we aimed to saturate them with elevated levels of GTP. To achieve this, we increased intracellular GTP levels by stimulating the nucleoside salvage pathway (Figure 5A) through supplementation of the growth medium with guanosine. ^37,38^ As a control we used supplementation with adenosine, which results in elevation of ATP levels. HEK293 cells were cultured for 48 hours in medium supplemented with guanosine, adenosine or DMSO serving as a vehicle control. Following this preconditioning, cells were transfected with 76 bp dsRNA bearing either 5′-pppA or 5′-pppG termini to assess the impact of altered purine nucleotide pools on dsRNA sensing and signaling. Following guanosine supplementation, transfection with both 5′-pppA and 5′-pppG 76 bp dsRNAs resulted in elevated IFN production compared to DMSO or adenosine-treated controls (Figure 5B-D). Importantly, the difference in IFN response, evaluated by measuring IRF3 phosphorylation and RIG-I levels, between 5′-pppA and 5′-pppG dsRNAs was substantially reduced in guanosine-enriched conditions (Figure 5B-D). These findings suggest that *in cellulo* elevated GTP levels enhance the RIG-I/IFN response and saturate GTP-binding proteins that normally bind 5′-pppG dsRNA and block RIG-I recognition through steric interference.

**Fig. 5.**
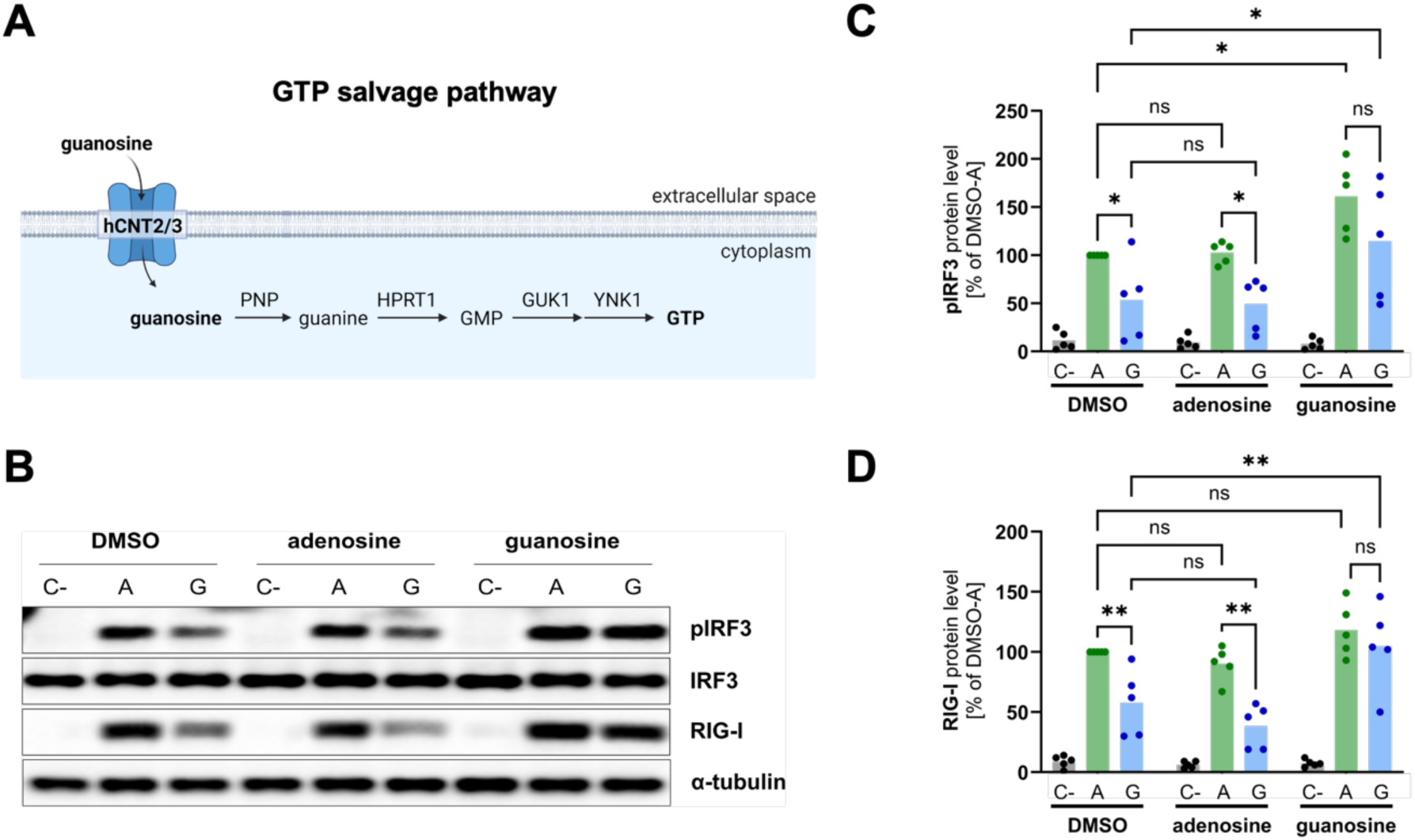
Increase of cellular GTP level through nucleoside salvage pathway minimizes differences in immune response to 5′-pppA and 5′-pppG dsRNAs. (**A**) Schematic presentation of the GTP salvage pathway. (**B**) Western blot analysis of IRF3 phosphorylation and RIG-I protein levels in HEK293 cells treated with guanosine, adenosine (control), or DMSO (vehicle) and then transfected with lipofectamine alone (C−), 5′-pppA or 5′-pppG IVT 76 bp dsRNAs. (**C, D**) Densitometric analysis of IRF3 phosphorylation (**C**) and RIG-I protein level (**D**) in HEK293 cells treated with guanosine, adenosine (control), or DMSO (vehicle) and then transfected with lipofectamine alone (C−), 5′-pppA or 5′-pppG IVT 76 bp dsRNAs. The results presented are derived from five independent western blot analyses. Two-way ANOVA with Šídák’s multiple comparisons test was used for statistical analysis.

In summary, our study demonstrates that dsRNAs commencing with 5′-pppA trigger a more robust RIG-I/IFN response in comparison to their corresponding counterparts initiating with 5′-pppG. The observed phenomenon holds across various tested RNAs, cell lines as well as in mice and human cells. Our working hypothesis proposes that GTPases and GTP-binding proteins impede the activation of RIG-I by 5′-pppG RNAs within the cellular environment, thereby diminishing their immunogenicity (Figure 6).

**Fig. 6.**
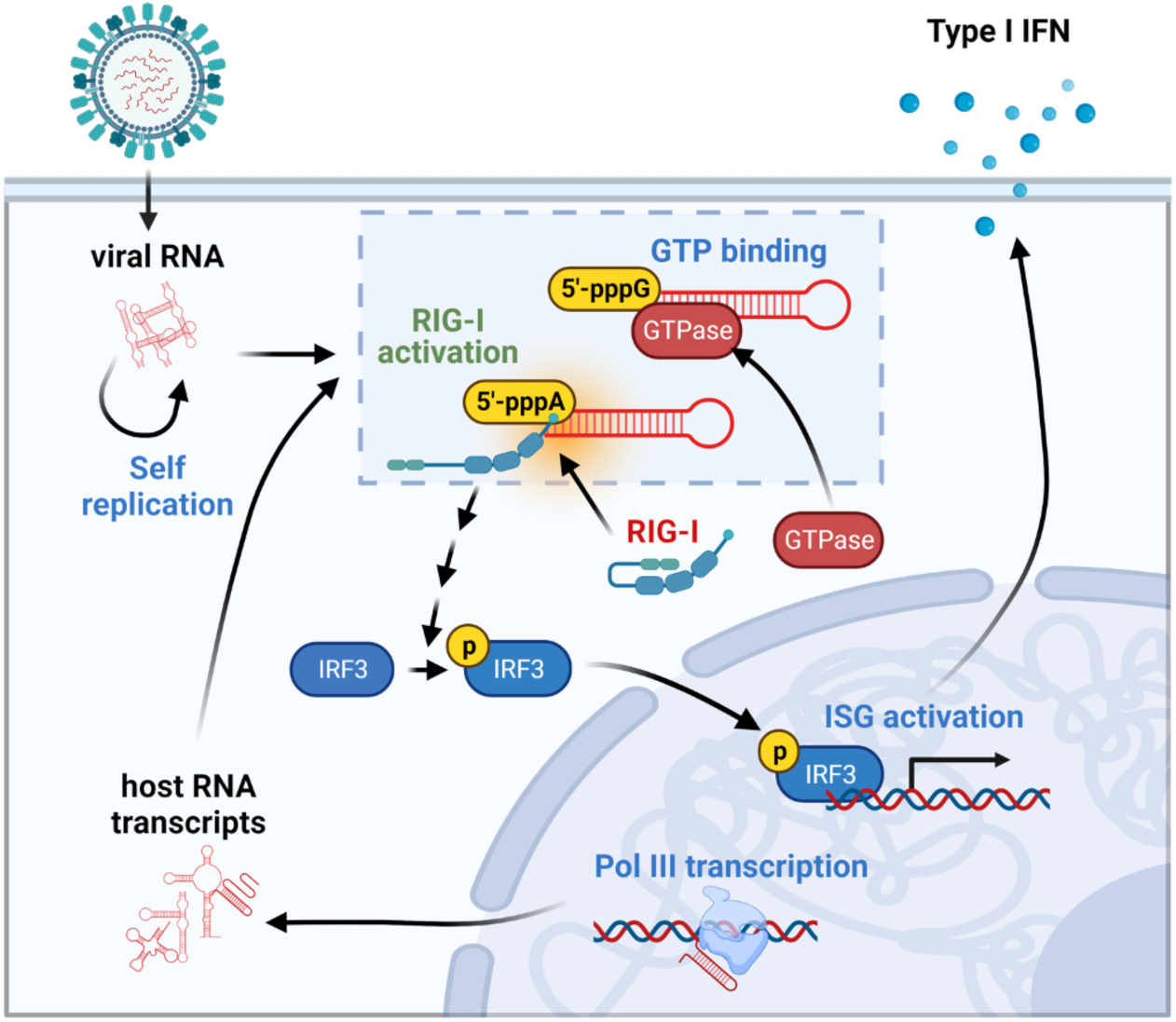
A model presenting a hypothesis of sequence-specific RNA sensing by RIG-I. Viral RNA, upon entry to the host cells, undergoes self-amplification. RIG-I may sense Pol III transcripts and viral RNAs starting with 5′-pppA. This recognition triggers a cascade of events, including IRF3 phosphorylation, ISG activation, and the subsequent production of type I IFN. RNAs containing a terminal nucleotide of 5′-pppG are identified by GTPases and GTP-binding proteins, which subsequently obstruct their recognition by the RIG-I/IFN pathway.

## Discussion

Viruses possess a range of conserved features, including uncapped 5′-ppp termini, unmethylated CpG-rich motifs, or dsRNAs. Although these features are typically absent or occluded in eukaryotic host RNAs, they function as targets for host innate immune system proteins, including RIG-I. ^39,40^ Under pathological conditions, host RNAs can trigger the RIG-I pathway. ^41,42^ To avoid erroneous activation of this pathway, host RNAs have evolved distinct molecular features and some characteristics that prevent recognition by the innate immune system. These include the presence of cap structures and nucleotide modifications on RNA polymerase II-derived mRNAs ^42^, specific cleavage patterns and chemical modifications on RNA polymerase I-transcribed rRNAs ^43^, and the clearance of 5′-ppp termini on RNA polymerase III transcripts, achieved through the activity of phosphatases such as triphosphatase dual-specificity phosphatase 11 (DUSP11). ^44^ Additionally, subcellular compartmentalization, as seen in the restricted localization of mitochondrial dsRNA, further limits unwanted immune detection. ^44^ Collectively, these adaptations ensure that host RNAs remain largely undetected by the innate immune system under physiological conditions.

Here we propose a new conceptual model underlying sequence-specific RNA recognition by RIG-I and the ensuing adequate activation of the type I interferon response (Figure 6). In this model, endogenous RNAs, representing most Pol III transcripts in higher eukaryotes, and certain highly pathogenic viruses initiate with 5′-pppG. This would make them relatively weak RIG-I agonists due to interference by GTP-binding proteins and GTPases. Notably, GTPases typically exhibit affinities for GTP in the nanomolar range, whereas known ATPases show affinities in the high micromolar range.^45,46^ More recent study indicates that GTP-binding affinities are approximately 3,000-fold higher than those of ATPases.^47^ This difference may explain why RNAs with 5′-pppG termini are more affected than those with 5′-pppA ends. This and additional mechanisms, such as dephosphorylation by DUSP11, ^44^ could contribute to evading detection by the RIG-I signaling pathway. Conversely, RNAs commencing with 5′-pppA, a characteristic of more prevalent viruses like IAV or HCV, exhibit greater potential for RIG-I/IFN pathway activation, which drives them to evolve mechanisms to silence or evade immune response. In the case of IAV, its NS1 protein has multiple functions that interfere with the host immune system ^48^, while in HCV, subtypes carrying 5′-pppA but not 5′-pppG are capped by FAD.^21^ Although some reports suggest that FAD might not be hindering RIG-I activity.^49^ Additionally, IAV is known to use a 5′-cap-snatching strategy to hide its transcripts from innate sensors and secure efficient cap-dependant translation. ^50^ The results presented here demonstrate that IVT-produced or chemically synthesised dsRNAs initiating with 5′-pppA are more immunogenic than those starting with 5′-pppG. Similar differences for short dsRNAs have been observed before but remained unexplored.^10^ Substantiating these findings is the observation that the Pol III transcript Y5 RNA, bearing 5′-pppA, is expressed at markedly lower levels compared to other Y RNAs initiating with 5′-pppG (Supplementary Figure S1) and is retained in the nucleus.^27^ Taken together, we speculate that the preservation of more immunogenic 5′-pppA termini in viruses may be influenced by constraints imposed by promoter and polymerase functions. Conversely, elevated IFN signaling could help maintain a balance in viral replication, allowing propagation without killing the host.

The biggest difference in the induction of the I interferon upon transfection with 5′-pppA or 5′-pppG RNAs occurred at very low concentrations. This could represent the early stages of virus infection which are crucial to mount an effective cellular response. ^51^ Similarly, low levels of immunogenic viral RNAs derived from West Nile virus and generated during the viral replication cycle have been shown to activate the RIG-I response.^52^ At the highest RNA concentrations tested (100 ng/ml for IVT 76 bp dsRNA and 1 µg/ml for synthetic 24 bp dsRNA), no significant difference was observed between the 5′-pppA and 5′-pppG variants. This implies an involvement of endogenous factors that may become saturated at very high concentrations of transfected RNAs. Intriguingly, Ren *et al.* reported no difference in IFN-β induction between dsRNAs starting with 5′-pppA or 5′-pppG. ^53^ The apparent discrepancy with our study may be attributed to different concentrations tested and the distinct dsRNA substrates examined. Ren *et al*. conducted their experiments using fully complementary dsRNAs with two exposed ends at a concentration of 2 µg/ml. Given that their RNAs had a length of 24 nt with 10 bp, the corresponding molar concentration was approximately 259 nM. In contrast, our IVT-produced 76 bp dsRNA, with a length of 76 nt, exhibited optimal results at a concentration of 20 pM (equivalent to 1 ng/ml) and, similarly to the high concentrations tested by Ren *et al*., showed no discernible difference in IFN induction at concentrations of 2 nM (equivalent to 100 ng/ml) and higher. Moreover, the minimal length documented thus far for the RIG-I agonist was 23 bp ^17^ and 10 bp. ^54^ We postulate that, beyond base pairing considerations, the overall length of the ligand is pivotal for RIG-I recognition. Supporting a preference for longer RNAs, Velthuis *et al.* demonstrated that mini viral RNAs ranging from 56 to 125 nucleotides, derived from the IAV, triggered the highest IFN-β promoter activity. ^16^

To investigate the mechanisms underlying the differential activation of the RIG-I/IFN type I signaling pathway by 5′-pppA *vs.* 5′-pppG dsRNAs, we employed a combination of biochemical and cellular methods. One of the primary advantages of biochemical assays is their ability to be conducted under defined, isolated conditions, enabling precise characterization of molecular interactions through the quantification of direct binding parameters. ^55,56^ Using a biochemical approach, we observed that RIG-I exhibits a marginally higher binding affinity for 5′-pppA dsRNA than for 5′-pppG dsRNA. Previous work has shown a correlation between RNA binding affinity to RIG-I and the capacity of RNA agonists to induce the RIG-I/IFN signaling pathway. ^57^ These findings support the idea that the identity of the 5′ nucleotide can influence the immunogenic potential of dsRNA ligands. In contrast, RIG-I activity, assessed by ATP consumption, showed no significant difference upon binding to 5′-pppA or 5′-pppG dsRNA (Figure 3). However, other studies have demonstrated that the ATPase activity of RIG-I does not fully reflect its functional activation. This is because RIG-I activation primarily depends on ATP binding, which drives essential conformational changes, rather than on ATP hydrolysis itself. ^55,56^ Thus, biochemical assays did not fully explain the observed difference between 5′-pppA and 5′-pppG dsRNA immunogenicity.

Since biochemical methods did not provide a definitive explanation for the mechanism underlying the differential activation of the RIG-I/IFN pathway by 5′-pppA and 5′-pppG dsRNAs, we investigated whether additional cellular factors might contribute to the observed phenomenon. Here, we employed an RNA pull-down assay combined with mass spectrometry, enabling the identification of proteins that specifically bind to the RNA of interest. We demonstrated that even a single nucleotide difference in the dsRNA sequence can alter the protein-binding profile, leading to the enrichment of distinct protein groups. Interestingly, when using 5′-pppA dsRNA as bait, we observed the enrichment of proteins typically associated with ssRNA, despite the dsRNA being immobilized on the resin. This finding suggests that A-U base pairs, being less thermodynamically stable, may allow partial or full unwinding of the duplex in the presence of helicases or other RNA-binding proteins in the lysate. Consequently, these transiently exposed ssRNA regions may become accessible to ssRNA-binding proteins. In comparison, Gene Ontology analysis of the 5′-pppG dsRNA pull-down revealed nine proteins classified as GTPases or GTP-binding proteins. Among them, a particularly novel finding is the identification of NUDT16, a known decapping enzyme that catalyzes the cleavage of the 5′ cap structure in snoRNAs and mRNAs. ^58^ Here, NUDT16 was found to bind to uncapped dsRNA, a previously unreported interaction. Furthermore, our results show that its binding to short dsRNA is strictly dependent on the presence of the 5′-pppG moiety and can be disrupted by blocking the GTP-binding pocket of NUDT16 with a non-hydrolyzable GTP analog. Although our experiment does not establish whether NUDT16 binds directly to dsRNA or as part of a larger protein complex, its interaction with uncapped RNA represents a novel discovery. Importantly, while NUDT16 has not been directly implicated in the activation or inhibition of the RIG-I/IFN innate immune signaling pathway, emerging evidence suggests that it contributes to innate immunity by regulating the turnover of inflammatory mRNAs and maintaining genomic integrity in immune cells. Its expression is upregulated under inflammatory conditions, such as sepsis, indicating a potential modulatory role in controlling the intensity of the innate immune response. ^59,60^

Of particular significance is the observed upregulation of the RIG-I/IFN signaling pathway in response to 5′-pppG dsRNA upon stimulation of the GTP salvage pathway. Combined with the finding that GTPase interaction with dsRNA is disrupted when its GTP-binding pocket is occupied, these results demonstrate that elevated GTP levels might relieve the inhibitory interaction between GTPases and 5′-pppG dsRNA. Notably, many viral infections lead to host mRNA degradation, resulting in increased levels of nucleotide precursors via the salvage pathway, which viruses exploit to support their replication. ^61^ According to our findings, this increase of nucleotide levels, including GTP, may also benefit the host by enhancing activation of the RIG-I/IFN pathway. Endogenous RNA polymerase III transcripts bearing 5′-pppG ends, such as Y-RNAs, may act as stoichiometric immunostimulatory ligands under these conditions, thereby contributing to the innate immune response during viral infection. In addition to RNA derived from RNA viruses, DNA viruses can also be a source of RNAs recognized by RIG-I. Consequently, many viruses may harbor factors that inhibit RIG-I activity. ^17^ Notably, RNAs from these DNA viruses (e.g., Epstein-Barr virus (EBV)) also produce RNAs with 5′-pppA termini, which could be highly immunogenic. ^62^ It is also worth noting that EBV infection is linked to autoimmune diseases such as systemic lupus erythematosus, ^63^ multiple sclerosis, ^64^ and rheumatoid arthritis. ^65^ It is suggested that EBV may contribute to the onset or exacerbation of autoimmune disorders. Despite these insights, the precise role of such RNAs in autoimmune diseases remains unclear, warranting further investigation of the complex interplay between the virus and the host’s immune system.

### Summary

In summary, our results suggest that accurate recognition of uncapped dsRNA by RIG-I in the cells may rely on the presence of a specific terminal 5′-ppp nucleotide and the surrounding cellular protein environment, with RNAs containing 5′-pppA being more immunogenic than those with 5′-pppG. This novel aspect of sequence-specific signaling in the RIG-I/IFN pathway will be important for advancing knowledge and designing strategies to target pathogenic viruses and host RNAs involved in autoimmunity.

### Limitations of the study

While we demonstrated that dsRNAs containing 5′-pppA are more immunogenic than dsRNAs with 5′-pppG, we could not have shown it for pol III transcripts transcribed *in vitro* or for RNA viruses. This is largely due to the IVT bias towards producing more immunogenic dsRNA side products when initiation occurs with 5′-pppA, as well as evolutionary and mechanistic pressures acting on viral promoter sequences. To address this limitation, future studies should focus on endogenously modified pol III transcripts or more extensive viral mutagenesis. However, any viral RNA modifications that enhance robustness through innate immune avoidance should be considered with caution.

## Materials and Methods

### IVT reaction

RNA transcripts (Supplementary Table S2) were prepared using the IVT reaction. IVT 76 bp dsRNA was derived by truncating 76 nucleotides from the 8^th^ segment of the IAV genome (GenBank: NC_002020.1).

The transcription template was first amplified using the High-Fidelity Phusion DNA Polymerase (Thermo #F530L) and primers (Supplementary Table S3) appending T7 Class III (TAATACGACTCACTATA) or ϕ2.5 Class II (TAATACGACTCACTATT) promoter sequence to produce 5′-pppA RNA or 5′-pppG RNAs, respectively. ^66^ Subsequent IVT reaction producing 5′-triphosphorylated RNAs was performed with NxGen T7 RNA Polymerase (Biosearch Technologies #30223-1).

Then RNA was precipitated with 3 M sodium acetate pH 5.1 and 100% EtOH, washed with 100% EtOH, and resuspended in UP water. The RNA in 2× loading buffer (7 M urea, bromophenol blue, and xylene cyanol) was then run on a denaturing polyacrylamide gel (10% polyacrylamide, 7.5 M urea in 1× TBE) for 2 h. Bands were stained with Stains-all (Sigma-Aldrich #E9379) and excised with individual scalpel blades. Then RNA was extracted (0.3 M sodium acetate pH 5.2; 0.5 mM EDTA; 0.1% SDS) and precipitated again. Washed with 100% EtOH and resuspended in UP water. RNA concentrations were assessed with NanoDrop. RNA was aliquoted and stored at −80°C. PAGE/urea gel electrophoresis was repeated to confirm that no extra bands were observed in the final RNA preparation. Dephosphorylated RNA was produced with FastAP enzyme (Thermo #EF0654) and cleaned using column purification (Invitek Invisorb Spin Virus RNA Mini Kit).

Double-stranded RNAs were prepared by mixing a 1:1 molar ratio of sense and antisense strands in 1× RNA refolding buffer (100 mM Tris-HCl pH 7.5, 100 mM NaCl, and 5 mM MgCl_2_). To obtain proper duplex formation, dsRNA mixture was heated up to +80°C for 10 min and then cooled down at room temperature for 30 min.

The 3p-hpRNA at a concentration of 100 ng/ml (InvivoGen #tlrl-hprna) or no RNA were used as a positive and negative controls, respectively.

### Chemical RNA synthesis

Standard solid-phase oligonucleotide synthesis was performed on an Äkta Oligopilot 10 Plus from GE Healthcare in a 4 µmol scale using 2ʹOTBDMS protected phosphoramidites and the 5ʹtrityl-off modus. Commercially available phosphoramidites, CPG materials and reagents were used, purchased from Sigma Aldrich, Avantor, TCI, and Glen Research.

The triphosphorylation reaction was performed subsequently after solid-phase synthesis with an automated machine adapted version of Goldeck et al. on an Äkta Oligopilot 10 Plus from GE Healthcare. ^67^ For deprotection, the CPG-bound oligonucleotide was incubated with 1.2 ml 40% methylamine for 45 min at +45°C. The material was diluted in 9 ml DMSO and then 900 µl glycolic acid was added. The 2ʹ deprotection was carried out using 1 M NH_4_F in DMSO for 8 h at room temperature. The reaction was quenched with 1 ml of 100 mM triethylammonium bicarbonate (TEAB) and passed through a Dowex Et_3_NH^+^ column for ion-exchange prior to HPLC purification. RP-HPLC purification was performed on a 15RPC Tricorn 10/300 with a linear gradient of 0-80% B in 30 CV at a flow rate of 2 ml/min. Buffer A was 100 mM TEAB and buffer B was 100 mM TEAB in 80% methanol (MeOH). For removal of the 5ʹdecyl-NH-tag, the product fractions were collected, evaporated and desalted by repeated co-evaporation with MeOH. The residue was redissolved in 400 μl deprotection buffer (100 mM acetic acid, TEMED, pH 3.8) and heated to +60°C for 70 min. After cooling the reaction mixtures on ice, 24 μl of 5 M NaCl and 1.2 ml ethanol were added to precipitate the oligonucleotide. The pure product was pelleted by centrifugation, washed with ethanol, dried under vacuum, and dissolved in pure water.

Purity and identity were confirmed by LC-MS analysis which was performed using an ACQUITY-UPLC system (Waters) coupled to an Xevo TQ-S Quadrupole (Waters) equipped with an electrospray source operating in negative ionization mode. All samples were chromatographed on an ACQUITY UPLC BEH C18 column (2.1 × 50 mm; 1.8 μm particle size) at +60°C column temperature. Separation of the analytes was achieved using a gradient of 16.6 mM triethylamine (TEA), 100 mM hexafluoroisopropanol (HFIP) and 10% methanol as eluent A and 16.6 mM TEA, 100 mM HFIP and 95% MeOH as eluent B with a flow rate of 0.3 ml/min. Samples were prepared in 200 mM TEAA and 20 mM EDTA. Segmented gradient for LC-MS analysis is presented in the Supplementary Table S4.

### Native polyacrylamide gel electrophoresis

For native polyacrylamide gel electrophoresis, a 10× native gel loading buffer (1 mM Tris-HCl pH 7.5, 5% glycerol, 0.001% bromophenol blue) was added to the samples and mixed. The mixtures were overlaid on a nondenaturing 12% polyacrylamide gel (prerun step was performed at 8 W for 45 min at +4°C). After electrophoresis at 8 W for 120 min at +4°C, the gels were stained with SYBR Gold nucleic acid gel stain (Invitrogen #S11494). Imaging was performed with a Chemidoc MP (Bio-Rad Laboratories).

### Cell cultures

Murine bone marrow cells were obtained from C57BL6J/Rj mice as previously described with modification. ^68^ Bone marrow cells were cultured with 20% of L929 conditioned medium for 7 days to generate BMDMs.^69^ Human HEK293, HEK-Blue IFN-α/β (InvivoGen #hkb-ifnab) cells, and murine BMDM cells were maintained in Dulbecco’s Modified Eagle’s Medium (DMEM; Gibco #32430) supplemented with 10% fetal bovine serum (Gibco #10270-106). HEK-Blue IFN-α/β cells were cultured with 100 μg/ml Zeocine (InvivoGen #ant-zn-1) and 30 μg/ml Blasticidine (InvivoGen #ant-bl-05). Primary cell cultures (BMDMs) were kept in presence of 1× Penicillin-Streptomycin (Gibco #15140122) and 0.1 mg/ml Gentamicin (Gibco #15710064). Cells were cultured at +37°C in a 5% CO_2_ humidified incubator.

### Transfection of RNAs in cell cultures

Cells were seeded in a 12-well plate at a concentration of 0.3-0.7×10^6^ cells per well and incubated for 24-48 hours. RNA was mixed with Lipofectamine: first, RNA was prediluted in 125 μl of OptiMEM (Gibco #11058), second, 2 μl of Lipofectamine 2000 (InvitroGen #11668) prediluted in 125 μl of OptiMEM was added, third, mix was incubated at RT for 30 min, fourth, 750 μl of cell culture medium was added. The obtained mix of RNA with Lipofectamine in medium was added to cells and incubated for 24 hours. Supernatants and cell lysates in modified Roeder D Buffer^70^ (20% (w/v) glycerol, 100 mM KCl, 0.2 mM EDTA; 100 mM Tris-HCl pH 8.0; 0.2 mM PMSF and 0.5 mM DTT) were collected and processed either for type I IFN assay or western blot.

### HEK-Blue IFN type I assay

Supernatants from HEK293 cells were processed with the HEK-Blue IFN type I assay in quadruplicates. 20 μl of supernatants (undiluted or prediluted 1:7) were added to 50,000 HEK-Blue IFN-α/β cells per well in a 96-well plate. A standard curve was generated in parallel by serial dilutions of recombinant IFN-β standard in DMEM (R&D Systems #8499-IF-010). After overnight incubation (24 h), 20 μl of supernatant was mixed with 180 μl of the working solution of the QUANTI-Blue reagent (InvivoGen #rep-qbs) and incubated at +37°C for 0.1-3 h. Absorbance was measured at 654 nm using a Tecan Sunrise absorbance microplate reader. Blank values were subtracted from all wells and the four-parameter logistic (4PL) standard curves were fitted to provide semiquantitative analyses of the IFN concentrations produced in the RNA-transfected cells. ^71,72^

### Western blot analysis

Cell monolayers were washed once with ice-cold 1× DPBS (Gibco #14190) and resuspended in 50 μl of a Roeder D buffer. After vortexing, cells were sonicated with Diagenode’s Bioruptor Pico sonication device (10 cycles of 30 s ON/30 s OFF at +4 C) and centrifuged (10000×g, 5 min, +4°C). Supernatants were moved to pre-chilled low protein binding tubes and, after quantifying protein concentration with NanoDrop, stored at −80°C. Twenty to hundred μg of protein extract was resolved using a 10% gel. Proteins were transferred to nitrocellulose membranes (Amersham #10600007) using a wet transfer apparatus. Membranes were blocked with 1% Western Blocking Reagent (WBR; Roche #11921681001) for 1 hour at room temperature. Membranes stained for pIRF3 were blocked with 5% BSA in TBST buffer. The blocked membranes were incubated with primary antibodies diluted in 0.5% WBR in TBST overnight at +4°C (Supplementary Table S5), washed three times with TBST, incubated with secondary antibodies diluted in 0.5% WBR in TBST for 1 hour at room temperature (Polyclonal Goat Anti-Rabbit Immunoglobulins/HRP, Agilent Dako #P0448, 1/2000), washed three times with TBST. For the chemiluminescence reaction, peroxide ECL reagents (Bio-Rad #170-5061) were applied to each membrane and then, the membranes were visualized and results quantified using an iBright 1500 Imaging System (Thermo).

### RNA extraction and qRT-PCR

RNA was extracted with Trizol (Invitrogen #15596018), and one-step qRT-PCR reactions were conducted on the Roche LightCycler 96 System. SYBRgreen-based GoTaq 1-Step qRT-PCR System (Promega #A6020) and LightCycler 96 software were used for RNA quantification. The reaction mixture (20 μl) consisted of 10 μl of 2× MasterMix, 0.4 μl of Reverse Transcription Mix, 0.2 μl of each primer diluted to 20 µM concentration, 7.2 μl of nuclease-free water and 2 μl (100 ng) of RNA template. A reverse transcription step of 15 min at +37°C and an enzyme activation step of 10 min at +95°C were followed by 40 amplification cycles (15 s at +95°C and 60 s at +60°C). The primers used are listed in Supplementary Table S6.

### Fluorescence RNA preparation

Fluorescent IVT dsRNA was constituted of sense strand RNA labelled on the 3′-end with fluorescein and unmodified antisense strand, both IVT derived. RNA 3′-end labelling was conducted as described earlier by Qiu *et al*.^73^ Briefly, 0.25 nmol of RNA was oxidized with 2.5 nmol of sodium periodate in RNA labelling buffer (0.25 M sodium acetate, pH 5.6), incubating in the dark for 90 min at room temperature. Then, 5 nmol of sodium sulphate was added, followed by 15 min incubation at room temperature. For labelling, 7.5 nmol of fluorescein-5-thiosemicarbazide was added into the mixture and incubated for 3 hours at +37°C. RNA was precipitated by adding 1/10 volume of 8 M LiCl and 2.5 volume of 100% ethanol and incubating on dry ice for 30 min (alternatively, overnight incubation in −20°C), followed by centrifugation at 16000×g at +4°C for 20 min. The RNA pellet was washed twice with 75% ethanol, resuspended in nuclease-free water and stored in −80°C.

Synthetic 24 bp dsRNA was constituted of sense strand unlabelled RNA and antisense strand RNA with 5′-end 6-FAM (6-carboxyfluorescein) modification prepared by IDT.

## Electrophoretic mobility shift assay (EMSA)

RNA-protein binding reaction was carried out in a 1× binding buffer containing 50 mM Tris-HCl pH 7.5, 2.5 mM MgCl_2_, 50 mM NaCl, 0.01% Tween-20 and 5 mM DTT. Serial dilutions of recombinant RIG-I protein were mixed with 0.5 nM fluorescently labelled dsRNA and 20 μM tRNA (Thermo #20159) as a non-specific competitor. Reactions were equilibrated for 30 minutes at room temperature, then mixed with 10× native gel loading buffer (1 mM Tris-HCl pH 7.5, 5% glycerol, 0.001% bromophenol blue) and loaded onto a 5% non-denaturing polyacrylamide gel. A pre-run was performed at 8 W for 45 minutes at +4°C. Free dsRNA and RNA-protein complexes were separated by electrophoresis at 8 W for 90 min at +4°C. Imaging was performed using a Typhoon laser scanner (Cytiva). The fraction of RNA bound to protein was calculated by densitometric analysis of the unbound fluorescent RNA band using the formula:

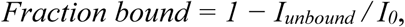

where: *I_unbound_* – the intensity of the unbound RNA band at a given RIG-I concentration; *I_0_* – the intensity of the unbound RNA band in the absence of RIG-I.

To estimate the apparent dissociation constant (K_d_), a 4PL model was fitted to densitometric data.

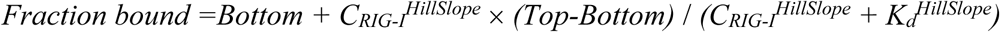

To compare K_d_ values across datasets, we used the entropy maximization principle and evaluated model fit using the Akaike Information Criterion corrected for small sample sizes (AICc). The probability that both datasets share a common Kd was calculated using the relative likelihood: Akaike’s Probability = 1 - 1 / (1 + e^ΔAICc/2^).

### RIG-I ATPase assay

The activity of recombinant RIG-I protein upon dsRNA binding under isolated conditions was assessed by measuring its ability to hydrolyze ATP to ADP. Similarly to the EMSA procedure, the ATP hydrolysis reaction was carried out in a 1× enzymatic reaction buffer. Briefly, a reaction mixture containing 1.3 nM RIG-I protein and 100 μM ATP, with varying concentrations of dsRNA, was prepared and incubated at +37°C for 60 minutes. ATP-to-ADP conversion was detected using Transcreener ADP^2^ FP Assay Kit (BellBrook Labs), following the manufacturer’s protocol. Fluorescence polarisation (FP) measurements were performed using a Tecan Infinite M1000 microplate reader.

To estimate the relative half-maximal effective concentration (EC_50_) of each RNA, a 4PL model was fitted to data representing the change in polarization across an RNA concentration gradient:

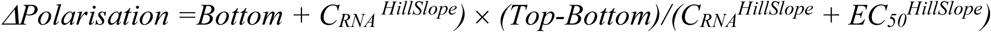

To compare EC_50_ values across datasets, Akaike’s methodology was applied as described for K_d_ estimation in the EMSA section.

### RNA pull-down

RNA pull-down assay was carried out to detect proteins binding to RNA immobilized on agarose beads. 250 pmol of *in vitro* transcribed and PAGE-purified sense strand RNA was treated with 100 mM sodium acetate and 5 mM sodium (meta)periodate in 200 μl of water and rotated for 1 h at room temperature in the dark. The RNA was precipitated by adding 600 μl of 100% ethanol and 15 μl of 3 M sodium acetate and incubating on dry ice for 30 min, followed by centrifugation at 16000×g, +4°C for 20 min. The RNA pellet was washed with 70% ethanol, followed by 5 min centrifugation at 16000×g. Then 250 pmol of antisense strand RNA in 50 μl 1× RNA refolding buffer (100 mM Tris-HCl pH 7.5, 100 mM NaCl, and 5 mM MgCl_2_) was used to resuspend the sense strand RNA pellet. RNA mixture was incubated for 30 min at room temperature and then 450 μl of 100 mM sodium acetate pH 5.2 was added.

For one reaction, 250 μl of adipic acid dihydrazide-agarose beads (Sigma-Aldrich #A0802) were washed 3× with 100 mM sodium acetate, followed by centrifugation at 2,000×g, +4°C for 2 min, then mixed with 500 μl of the periodate-oxidized RNA and incubated overnight at +4°C in the dark with rotation. The RNA-beads were washed by mixing with 700 μl of 4 M KCl, rocked for 30 min at room temperature, centrifuged at 2,000×g for 5 min, washed 2× with 2 M KCl, washed 2× with Buffer G (20 mM Tris-HCl pH 7.5, 137 mM NaCl, 1 mM EDTA, 1% Triton X-100, 10% glycerol, 1.5 mM MgCl_2_, 1 mM DTT and 200 μM PMSF) and washed 1× with Roeder D followed by 2 min centrifugation at 2,000×g at room temperature. Control beads with no RNA attached were also prepared. 1 mg of total protein extract was added to RNA-beads. The mixture was supplemented with 1.5 M MgCl_2_, 25 mM creatine phosphate, 100 mM ATP, and 2.5 μl of RiboProtect Hu RNase Inhibitor (Blirt #RT35). Additionally, for GTP analog treated samples Non-hydrolyzable GTP Test Kit (Jena Bioscience #NK-102) containing 5 analogs – GTPαS, GpCpp, GppCp, GppNHp, GTPγS – was used. Each GTP analog was added at 0.1 mM; combining all five analogs therefore produced a total nucleotide concentration of 0.5 mM, which corresponds to the physiological GTP level in mammalian cells. ^74^ The mixture volume was adjusted to 650 μl with nuclease-free water. The RNA-beads-cell lysates mixtures were incubated at +37°C for 30 min with shaking. After 3× washes with Buffer G, the beads were mixed with 60 μl of 5× Sample Buffer. Proteins captured by RNA were denatured at +95°C for 10 min with shaking. 30 μl of the supernatant was loaded onto SDS-PAGE or NuPAGE gel and western blot or mass spectrometry analysis was performed to detect the proteins, respectively.

### Mass spectrometry

Proteins were separated on gel (NuPAGE Novex 4-12% Bis-Tris gel, Thermo) in NuPAGE buffer (MES) for 10 min and visualised using InstantBlue stain (Abcam). The stained gel band was excised and de-stained with 50 mM ammonium bicarbonate (Sigma Aldrich) followed by 100% acetonitrile (Sigma Aldrich). Then proteins were digested with trypsin, as previously described. ^75^ In brief, proteins were reduced in 10 mM DTT (Sigma Aldrich) for 30 min at +37°C and alkylated in 55 mM iodoacetamide (Sigma Aldrich) for 20 min at ambient temperature in the dark. They were then digested overnight at +37°C with 13 ng/μl trypsin (Pierce). Following digestion, samples were diluted with an equal volume of 0.1% TFA and pH was set to ∼2 with 10% TFA.

The equivalent of 50 ng of the digest was loaded on the Evotip using the standard producer protocol. We utilized Evosep coupled to a TimsTOF ULTRA (Bruker) mass spectrometer equipped with a Captive Spray II source (Bruker). The separation was carried out on maintained at +40°C column (Evosep EV1115). We employed the standard 60SPD Evosep method, applying a 21-minute gradient for a total sample-to-sample time of 24 minutes using Solvent A (0.1% formic acid) and Solvent B (0.1% formic acid in acetonitrile); Thermo Optima LCMS grade).

The dia-PASEF acquisition scheme was optimized for a cycle time estimate of 1.38 s. The window scheme was designed to cover most of the charge 2 precursor ions in the range m/z 391−1142 and 1/K_0_ 0.68−1.36, using 22 × 31 Th windows, with accumulation and ramp times of 0.1 s. The mass spectrometer was operated in the high-sensitivity mode (“low sample amount”).

The DIA-NN software platform ^76^ version 1.9.1 was used to process the raw files from label-free DIA and the search was conducted against the *Homo sapiens* reference proteome [UniProt Proeome ID #UP000005640]. Precursor ion generation was based on the chosen protein database (automatically generated spectral library) with deep learning-predictions for spectra, retention times, and ion mobilities. Digestion mode was set to specific with trypsin allowing a maximum of one missed cleavage. Carbamidomethylation of cysteine was set as fixed modification. Oxidation of methionine, and acetylation of the N-terminus were set as variable modifications. MS1 and MS2 mass accuracies were set to 10 ppm. The parameters for peptide length range, precursor charge range, precursor m/z range and fragment ion m/z range as well as other software parameters were used with their default values. The precursor FDR was set to 1%.

Protein intensities were quantified from peptide intensities with directLFQ Python package.^77^ Statistical analysis was performed with R.^78^ *Protti* R package ^79^ was used for quality control, data filtration, imputation of missing values, and statistical significance calculation using a moderated t-test based on the *limma* R/Bioconductor package ^80^. For functional validation, *p*-values were adjusted for multiple testing with the Benjamini-Hochberg correction. Data visualisation was performed using the *ggplot2* R package.^81^ GO enrichment analysis for Biological Processes and Molecular Functions was conducted on the set of enriched proteins using the *clusterProfiler* package in R.^82^

### Murine BMDMs-based mKate2/IFN-β activity assay, image acquisition and analysis with Opera Phenix

Murine BMDMs-based mKate2/IFN-β activity assay was performed as described previously.^35^ In short, bone marrow-derived macrophages were isolated from a genetically modified mouse expressing the fluorescent protein mKate2 in place of the IFN-β (the mKate2 encoding sequence was inserted at the start of the IFN-β coding region with added SV40 polyadenylation signals and a mutation disrupting protospacer-adjacent motif-) and plated in 96-well plates (Greiner Bio-One #655090) at a density of 10^5^ cells per well. BMDMs were then transfected with different variants of dsRNAs and imaged hourly using the PerkinElmer Opera Phenix High-Content Screening System. Activation of the IFN-β promoter in these cells triggers mKate2 expression instead of IFN-β, allowing single-cell visualisation of IFN pathway induction. To enumerate cells, images were first flat-field–corrected, then segmented on the mKate2 channel with PerkinElmer Harmony software (“*Method C*” from the “*Find Cells”* building block).

### Production of the recombinant RIG-I protein

Gene coding RIG-I/DDX58 cDNA [GeneID: 23586] was cloned in pET28-6xHis-SUMO vector (Thermo #K30001). Protein was expressed in *E. coli* BL21-RIL cells (Agilent #230245) cultivated in Lysogeny Broth medium overnight at +18°C after induction with 0.4 mM isopropyl β-D-1-thiogalactopyranoside at OD_600_ of 0.6-0.7. The harvested cells were pelleted by centrifugation at 4,000×g, +4°C for 20 minutes and resuspended in buffer A (50 mM HEPES pH 7.5, 500 mM NaCl, 10% glycerol (w/v), 5 mM β-mercaptoethanol, 20 mM imidazole and 40 mM L-arginine) supplemented with proteinase inhibitors, and sonicated. The lysate was then subjected to ultracentrifugation at 142, 000×g, +4°C for 40 min yielding a clear supernatant. The clarified lysate was applied to a 5 ml His-Trap HP column (Cytiva #17524802) pre-equilibrated with buffer A. The column was washed with buffer A, followed by a wash with the same buffer that included 60 mM imidazole. RIG-I protein was eluted with buffer A that contained 500 mM imidazole. To remove the 6xHis-SUMO tag, the fusion protein was digested with SUMO protease (Ulp1) and dialyzed overnight against buffer A at +4°C. The cleaved-off His-tagged SUMO protein was removed on a His-Trap HP column. For the final purification step, tag-free RIG-I from the flow-through was concentrated and applied to a size exclusion column Hiload 16/600 Superdex 200 (Cytiva #28989335), pre-equilibrated with buffer A without imidazole. The RIG-I protein fractions were pooled and concentrated using a 30 kDa Amicon Ultra filter (Millipore #UFC903008). Lastly, the concentration of RIG-I was measured with NanoDrop and the purified protein was stored at −80°C.

### Induction of nucleotide salvage pathway

To elevate the GTP level in the cells, HEK293 12-well cell cultures were supplemented with 100 µM guanosine (Sigma Aldrich #G6264). DMSO and 100 µM adenosine (Sigma Aldrich #A9251) treatments were utilized as control conditions. The cells were incubated for 48 h in cell culture incubator and then transfected with IVT 76 bp dsRNAs at a final concentration of 0.1 ng/ml. After 24 h incubation the cells were lysed in Roeder D lysis buffer and used for western blot analysis.

### Statistical analysis

All data are reported as mean ± standard deviation. Statistical analyses were performed using GraphPad Prism 10.5.0.

## Data Availability

The mass spectrometry proteomics data have been deposited in the ProteomeXchange Consortium via the PRIDE ^83^ partner repository with the dataset identifier PXD065769. Supplementary data available online.

## Acknowledgments

Figures 2A, 2B, 4A, 5A, 6, Supplementary Figures S4A and S5A were created with BioRender.com.

## Author contributions

G.M. conceived this study; J.S., M.W. and G.M. designed the experiments; J.S., M.W., I.T., Z.N., N.I., E.N., and C.W., performed experiments; J.S., M.W., I.T., Z.N., N.I., G.H., J.R. and G.M. analyzed data; J.S., M.W. and G.M. wrote the manuscript with input from other authors. All authors read and approved the final manuscript.

## Funding

Project financed under DIOSCURI, a programme initiated by the Max Planck Society, jointly managed with the National Science Centre in Poland, and mutually funded by Polish Ministry of Science and Higher Education and German Federal Ministry of Education and Research [2019/02/H/NZ1/00002 to G.M.]; Polish National Agency for Academic Exchange within Polish Returns Programme as well as National Science Centre [2021/01/1/NZ1/00001 to G.M.]. This work was financed by the statutory funding of the International Institute of Molecular and Cell Biology in Warsaw. This research was performed thanks to the IIMCB IN-MOL-CELL Infrastructure [RRID: SCR_021630] funded by the European Union – NextGenerationEU under the National Recovery and Resilience Plan. IN-MOL-CELL Infrastructure was also funded by the European Union under Horizon Europe (Project 101059801–RACE) and by the RACE-PRIME project carried out within the IRAP program of the Foundation for Polish Science co-financed by the European Union under the European Funds for Smart Economy 2021–2027 (FENG). J.R. was funded by the Deutsche Forschungsgemeinschaft (DFG, German Research Foundation) under Germany′s Excellence Strategy (EXC 2008–390540038–UniSysCat) and project 449713269. The Wellcome Centre for Cell Biology is supported by core funding from the Wellcome Trust (Grant number 203149). Funding for open access charge: DIOSCURI Centre. This work was supported by the Deutsche Forschungsgemeinschaft (DFG, German Research Foundation) under Germany’s Excellence Strategy (EXC 2151–390873048) (G.H.).

## Competing interests

The authors declare no competing interests.

## Notes

### Competing Interest Statement

The authors have declared no competing interest.

### Summary of Updates

The manuscript title was revised to emphasise the immunosuppressive role of 5-prime pppG RNAs through interactions with GTP binding proteins. New experimental evidence demonstrates that GTP binding proteins selectively bind to 5-prime pppG RNAs, leading to suppression of RIG I signalling. To address potential artefacts arising from in vitro transcription (IVT), the RNA duplex generation strategy was refined, following the approach described by Wolczyk et al. (2025, NAR, PMID 39704128). Synthetic and IVT RNAs were hybridised from individually purified strands to ensure comparable structural integrity. Additional data were generated to compare binding affinities. EMSA and ATPase assays using purified RIG-I protein revealed similar activation profiles, with modest differences in the binding affinities between 5-prime pppA and 5-prime pppG RNAs. The study also expanded the range of RNA variants tested, including synthetic 24 base pair duplex RNAs, pre-let-7a, and Y5 RNA derivatives, thereby broadening the spectrum of RNA ligands assessed for immunogenic potential. Several supplementary figures were substantially revised or newly included. These additions comprise western blot validations of RIG I activation, EMSA results, and kinetic data from BMDM reporter cell assays. Numerous main figures were also updated to include time course data, statistical evaluations, and fold change comparisons, thereby enhancing the quantitative rigour of the analysis. The author list and affiliations were updated to reflect changes in contribution. Nathalie Idlin, Justyna Jackiewicz, Christine Wuebben, and Gunther Hartmann were added as co-authors, while Agnieszka Bolembach, Nila Roy Choudhury, Tola Tame, Ceren Konuc, and Christos Spanos were removed due to the exclusion of their experimental results. University Hospital Bonn was added to the list of institutional affiliations. The manuscript structure was reorganised, particularly within the introduction and results sections, to better support the mechanistic narrative. Clarifying context and methodological detail were added throughout. Textual revisions were made to improve precision, with standardised terminology and clearer descriptions of RNA constructs and experimental procedures. These collective updates enhance the mechanistic insight, methodological accuracy, and experimental depth of the manuscript.

