## Supplementary for "5′-triphosphate guanosine RNAs recruit GTP-binding proteins to suppress RIG-I/IFN type I signaling"

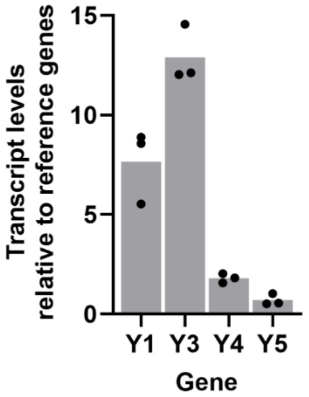


**Supplementary Figure S1. The levels of Y RNAs in HEK293 cells.** qRT-PCR for Y RNAs (primers listed in Supplementary Table S3) was done for HEK293 cells with three technical and three biological replicates. For normalization, we used the geometric mean of the expression levels for the three reference genes (*IPO8*, *GAPDH*, and *7SL*).

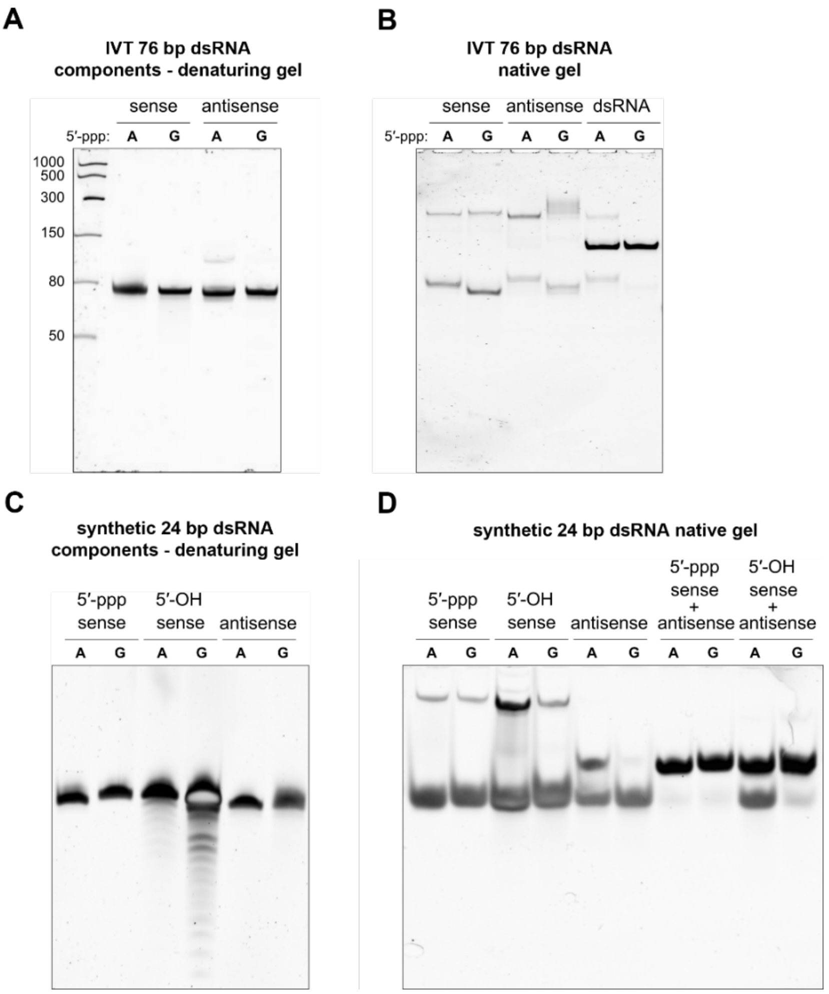


**Supplementary Figure S2. Quality control of IVT and chemically synthesised RNA products.** (**A, C**) PAGE/Urea analysis of IVT 76 bp (**A**) and synthetic 24 bp (**C**) dsRNA variants. (**B, D**) Native PAGE analysis showing individual single strands used for annealing and the resulting duplexes for the IVT 76 bp (**B**) and synthetic 24 bp (**D**) dsRNAs.

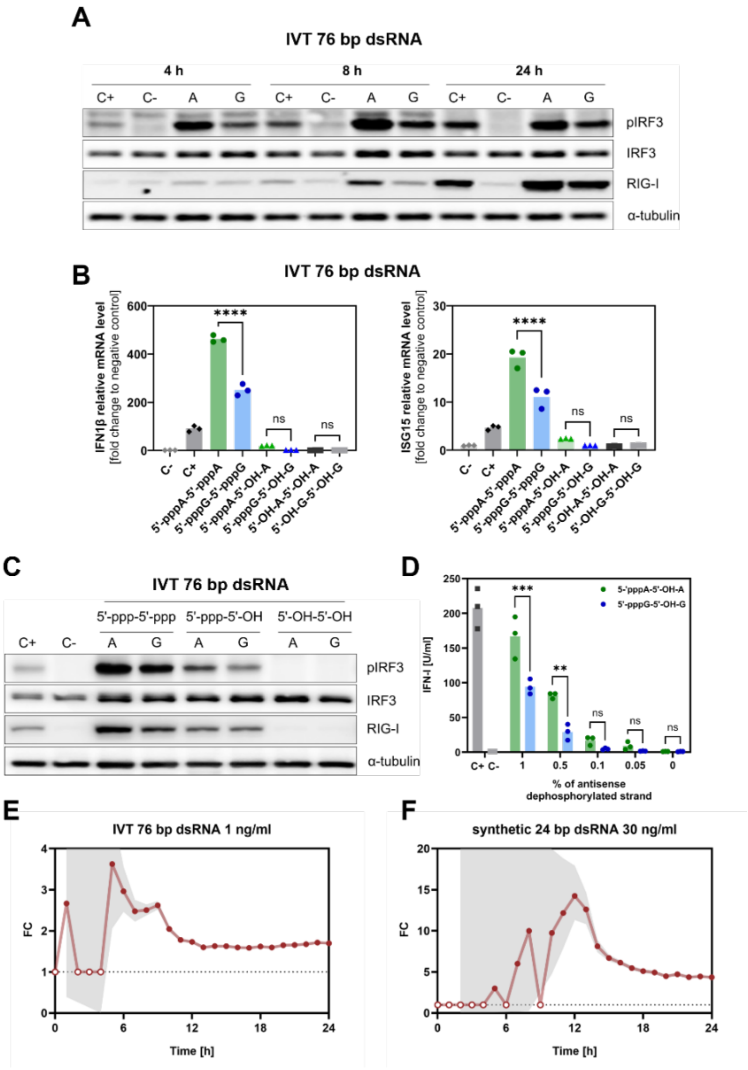


**Supplementary figure S3. 5′-pppA 76 bp dsRNA is more immunogenic than 5′-pppG 76 bp dsRNA.** (**A**) Analysis of IRF3 phosphorylation and RIG-I expression assessed by Western blot analysis in HEK293 cells after transfection with 1 ng/ml of 5′-pppA *vs.* 5′-pppG IVT 76 bp dsRNAs in 4, 8 and 24 h post transfection. (**B**) Analysis of *IFN1β* and *ISG15* relative mRNA levels in total RNA isolated from HEK293 cells after transfection with 1 ng/ml of 5′-pppA *vs.* 5′-pppG IVT 76 bp dsRNAs. Two-way ANOVA with Šídák's multiple comparisons test was used for statistical analysis (n=3). (**C**) IRF3 phosphorylation and RIG-I expression analysis in HEK293 cells after transfection with 1 ng/ml of 5′-pppA *vs.* 5′-pppG IVT 76 bp dsRNA variants (both strands triphosphorylated, only sense strand triphosphorylated, both strands dephosphorylated) 24 h post transfection. (**D**) Interferon activity measured using HEK Blue IFN type I assay in the supernatants collected from HEK293 cells 24 h after transfection with 3p-hpRNA (C+), lipofectamine (C−) and semisynthetic ligated 5′-pppA *vs.* 5′-pppG 76 nt RNA hybridized with different concentrations of fully complementary antisense dephosphorylated strand. Two-way ANOVA with Šídák's multiple comparisons test was used for statistical analysis (n=3). (**A-D**) The 3p-hpRNA at a concentration of 100 ng/ml (InvivoGen #tlrl-hprna) or lipofectamine were used as positive (C+) and negative (C−) controls, respectively. (**E-F**) Fold-change ratios of mKate2+ cell counts (FC=5′-pppA/5′-pppG) cells treated with either 76 bp IVT (**E**) or 24 bp synthetic dsRNAs (**F**). Grey band is a 95 % confidence interval obtained with the log-delta method. Time-points with either group mean was zero (ratio undefined) are marked by empty circles on the baseline (FC=1).

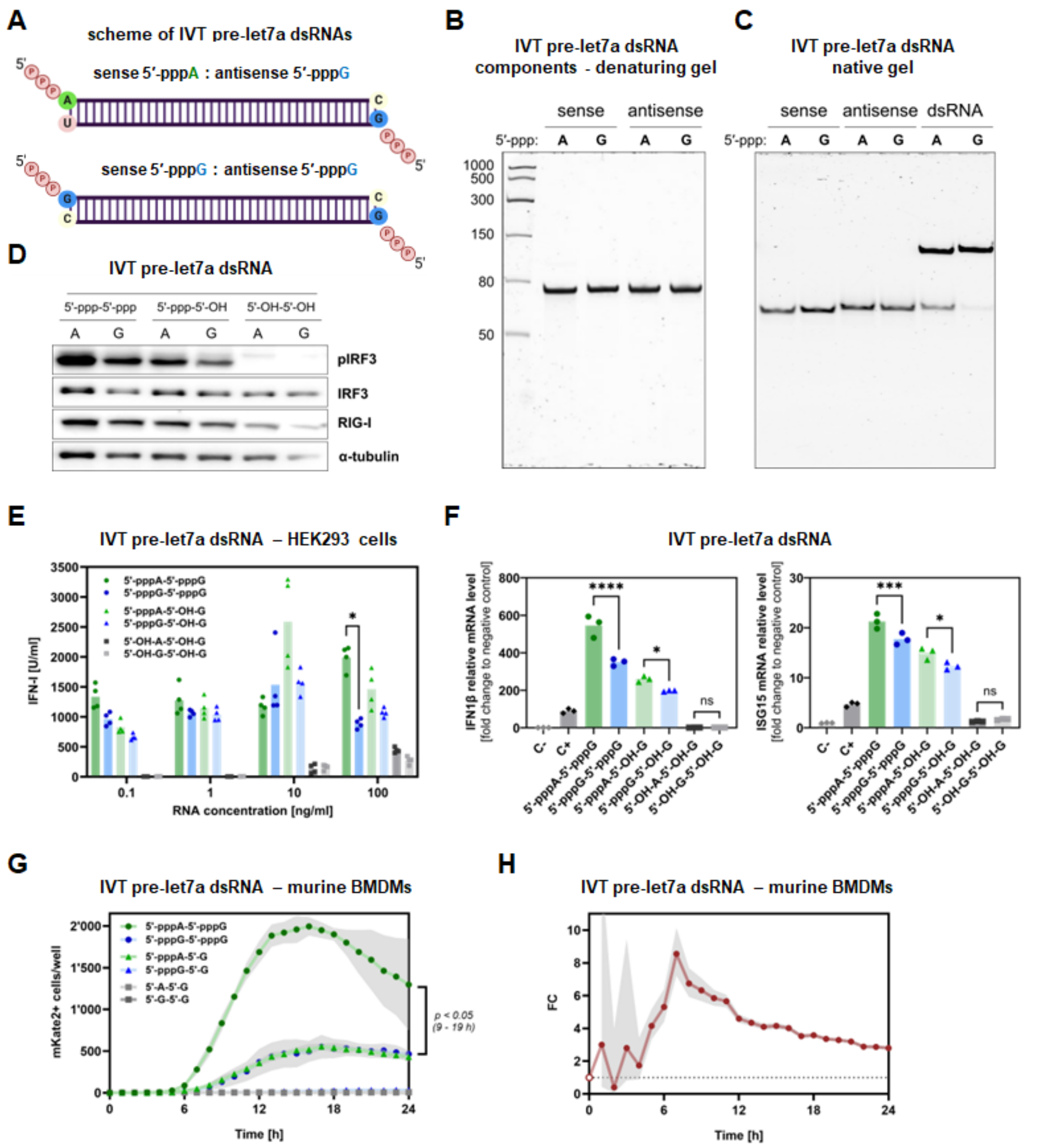


**Supplementary Figure S4. 5′-pppA pre-let7a dsRNA is more immunogenic than 5′-pppG pre-let7a dsRNA.** (**A**) Schematic representation of IVT-derived pre-let7a dsRNA variants differing at the 5′ terminal nucleotide (5′-pppA or 5′-pppG). (**B**) PAGE/Urea analysis of IVT-derived, single strand components of pre-let7a dsRNA variants. (**C**) Native PAGE analysis showing individual single strands of pre-let7a used for annealing and the resulting duplexes for IVT-derived pre-let7a dsRNA variants. (**D**) Analysis of IRF3 phosphorylation and RIG-I expression assessed by Western blot analysis in HEK293 cells after transfection with 1 ng/ml of IVT-derived pre-let7a dsRNAs with different 5′-terminal nucleotides 24 h post transfection. (**E**) Type I interferon production in HEK293 cells was assessed upon transfection with IVT-derived pre-let7a dsRNA variants with the HEK Blue IFN type I assay. Two-way ANOVA with Šídák's multiple comparisons test was used for statistical analysis (n=4). (**F**) Analysis of *IFN1β* and *ISG15* relative mRNA levels in total RNA isolated from HEK293 cells after transfection with 1 ng/ml of IVT-derived pre-let7a dsRNA variants. Two-way ANOVA with Šídák's multiple comparisons test was used for statistical analysis (n=3). The 3p-hpRNA at a concentration of 100 ng/ml (InvivoGen #tlrl-hprna) or lipofectamine were used as positive

(C+) and negative (C−) controls, respectively. (**G**) RIG-I/IFN activation in murine BMDMs upon transfection with IVT-derived pre-let7a dsRNA variants, shown as mKate2+ cell counts over 0–24 h post-transfection. Grey area represents standard deviation for n=5. Data were analyzed using repeated measures two-way ANOVA on log-transformed values [$\log_{10}(x + 1)$], with Geisser-Greenhouse correction and Šídák's multiple comparisons test. (**H**) Fold-change ratios of mKate2+ cell counts (FC=5′-pppA/5′-pppG) in murine BMDM cells treated with IVT-derived pre-let7a dsRNA variants differing at the 5′ terminal nucleotide (5′-pppA or 5′-pppG). Grey transparent band is a 95 % confidence interval obtained with the log-delta method. Time-points with either group mean was zero (ratio undefined) are marked by empty circles on the baseline (FC=1).

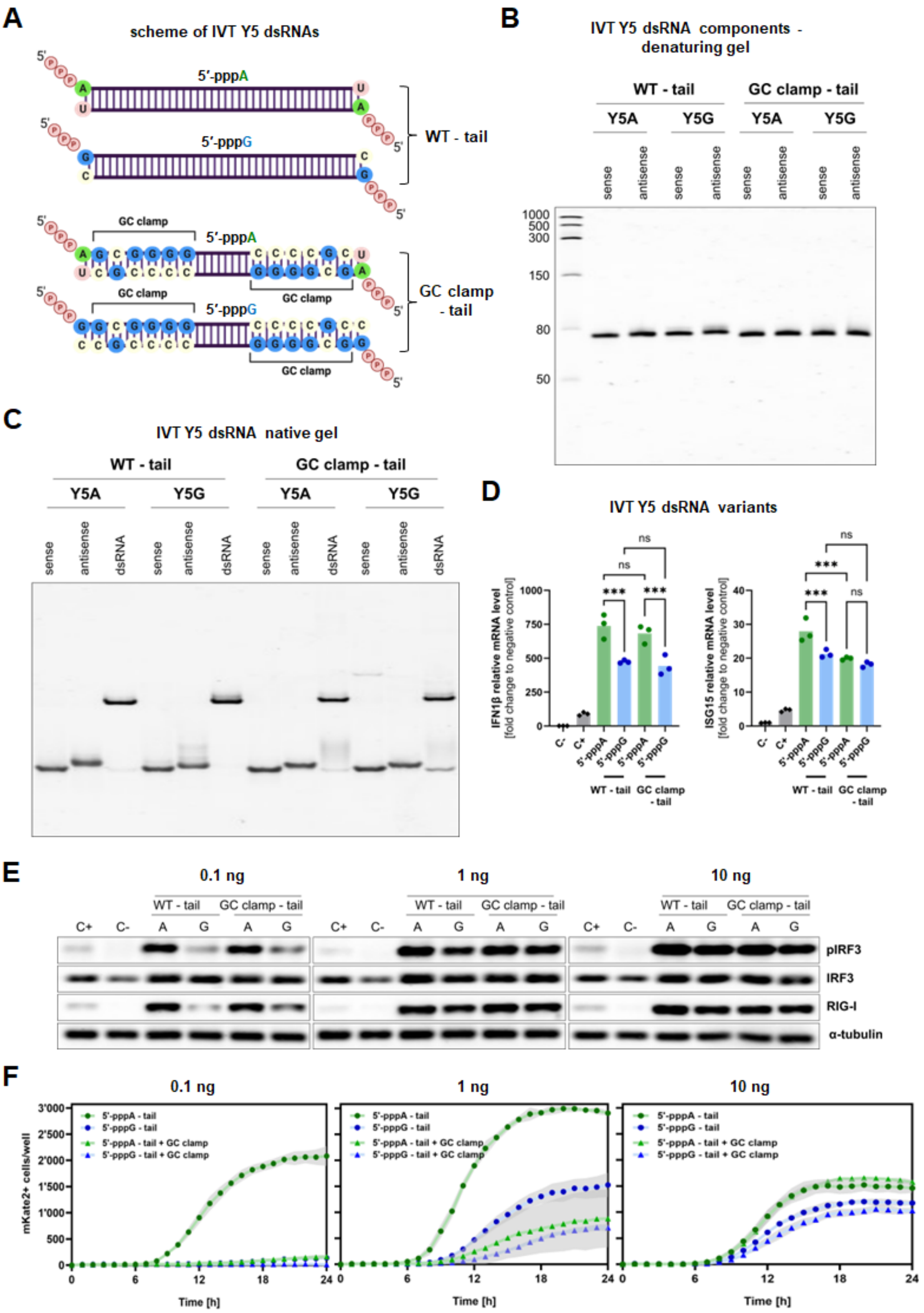
A
scheme of IVT Y5 dsRNAs
5′-pppA
5′-pppG
WT - tail
GC clamp
GC clamp - tail
B
IVT Y5 dsRNA components - denaturing gel
WT - tail
GC clamp - tail
Y5A
Y5G
sense
antisense
1000
500
300
150
80
50
C
IVT Y5 dsRNA native gel
dsRNA
D
IVT Y5 dsRNA variants
IFN1β relative mRNA level [fold change to negative control]
ISG15 relative mRNA level [fold change to negative control]
ns
***
C-
C+
5'-pppA
5'-pppG
E
0.1 ng
1 ng
10 ng
pIRF3
IRF3
RIG-I
α-tubulin
F
mKate2+ cells/well
Time [h]
5'-pppA - tail
5'-pppG - tail
5'-pppA - tail + GC clamp
5'-pppG - tail + GC clamp

G H

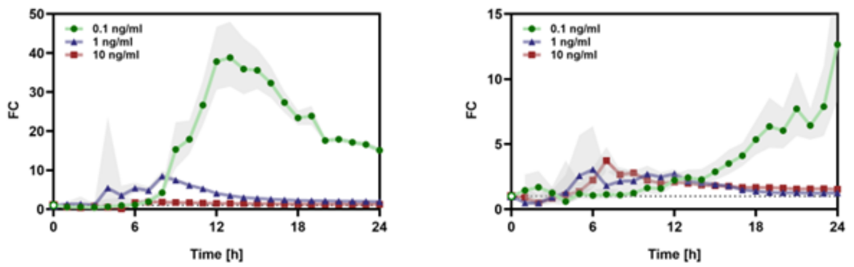


**Supplementary Figure S5. 5′-pppA Y5 dsRNA variants are more immunogenic than 5′-pppG Y5 dsRNA variants.** (**A**) Schematic representation of Y5 dsRNA variants. (**B**) PAGE/Urea analysis of IVT-derived, single strand components of Y5 dsRNA variants. (**C**) Native PAGE analysis showing individual single strands of Y5 RNAs used for annealing and the resulting duplexes for IVT-derived Y5 dsRNA variants. (**D**) Analysis of *IFN1β* and *ISG15* relative mRNA levels in total RNA isolated from HEK293 cells after transfection with 1 ng/ml of IVT-derived Y5 dsRNA variants. Two-way ANOVA with Šídák's multiple comparisons test was used for statistical analysis (n=3). (**E**) Analysis of IRF3 phosphorylation and RIG-I expression assessed by Western blot analysis in HEK293 cells after transfection with 10, 1 and 0.1 ng/ml of IVT-derived Y5 dsRNA variants 24 h post transfection. (**D** and **E**) The 3p-hpRNA at a concentration of 100 ng/ml (InvivoGen #tlrl-hprna) or lipofectamine were used as positive (C+) and negative (C−) controls, respectively. (**F**) RIG-I/IFN activation in murine BMDMs upon transfection with 10, 1 and 0.1 ng/ml of IVT-derived Y5 dsRNA variants, shown as mKate2+ cell counts over 0–24 h post-transfection. Grey area represents standard deviation for n=5. (**G-H**) Fold-change ratios (FC=5′-pppA/5′-pppG) for IVT-derived Y5 dsRNA variants without tail (**G**) and without tail and with GC clamp (**H**). Grey band is a 95 % confidence interval obtained with the log-delta method. Time-points with either group mean was zero (ratio undefined) are marked by empty symbols on the baseline (FC=1).

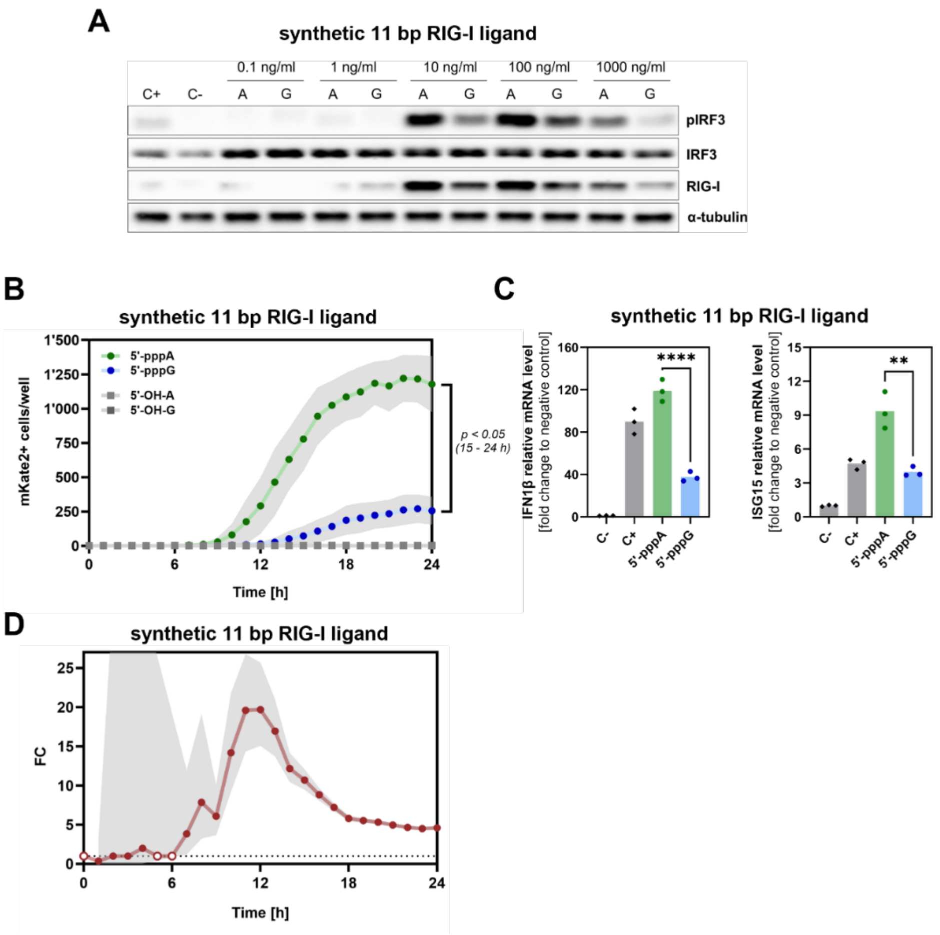


**Supplementary Figure S6. Synthetic 5′-pppA 11 bp RIG-I ligand is more immunogenic than synthetic 5′-pppG 11 bp RIG-I ligand.** (**A**) Analysis of IRF3 phosphorylation and RIG-I expression assessed by Western blot analysis in HEK293 cells after transfection with various concentrations of 5′-pppA *vs.* 5′-pppG short synthetic, partially double-stranded RIG-I agonist 24 h post transfection. (**B**) RIG-I/IFN activation in murine BMDMs upon transfection with tested synthetic dsRNAs, shown as mKate2$^{+}$ cell counts over 0–24 h post-transfection. Grey area represents standard deviation for n=5. Data were analyzed using repeated measures two-way ANOVA on log-transformed values [$\log_{10}(x+1)$], with Geisser-Greenhouse correction and Šídák's multiple comparisons test. (**D**) Fold-change ratios (FC=5′-pppA/5′-pppG) upon transfection of murine BMDM with synthetic dsRNAs. Grey band is a 95 % confidence interval obtained with the log-delta method. Time-points with either group mean was zero (ratio undefined) are marked by empty circles on the baseline (FC=1). (**C**) Analysis of *IFN1β* and *ISG15* relative mRNA levels in total RNA isolated from HEK293 cells after transfection with 1 ng/ml of IVT-derived Y5 dsRNA variants. Two-way ANOVA with Šídák's multiple comparisons test was used for statistical analysis (n=3). (**A** and **C**) The 3p-hpRNA at a concentration of 100 ng/ml (InvivoGen #tlrl-hprna) or no RNA were used as positive (C+) and negative (C−) controls, respectively.

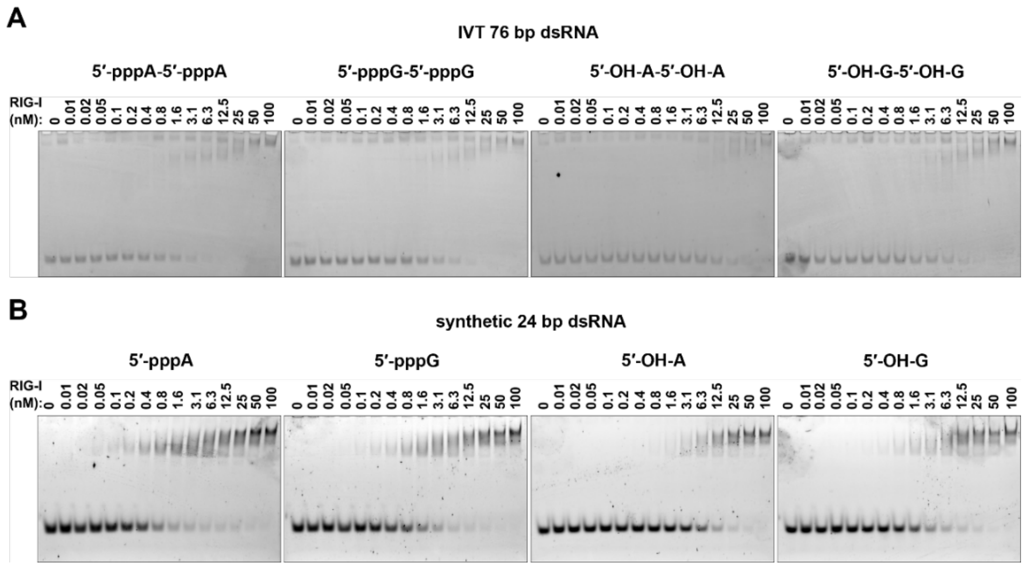


**Supplementary Figure S7. Electrophoretic mobility shift assay (EMSA) of recombinant RIG-I protein binding to the IVT 76 bp** (A) **and synthetic 24 bp** (B) **dsRNAs with different 5′-terminal nucleotides** (5′-pppA, 5′-pppG, 5′-OH-A, 5′-OH-G).

**Supplementary Table S1.** The dissociation constant ($K_d$) from EMSA and the half-maximal effective concentration ($EC_{50}$) for ATPase activity were compared across tested dsRNA variants. The entropy maximization principle was applied, and model fit was evaluated using the Akaike Information Criterion corrected for small sample sizes (AICc). Akaike probability threshold of 0.05 was used to determine statistical significance.

| RNA variant | Terminal Nucleotide Comparison | EMSA | | RIG-I Activity Assay | |
|---|---|---|---|---|---|
| | | Akaike's Probability | $K_d$ sigificantly different | Akaike's Probability | $EC_{50}$ sigificantly different |
| IVT 76 bp dsRNA | **5′-pppA vs 5′-pppG** | < 0.001 | Yes | 0.71 | No |
| | **5′-pppA vs 5′-OH-A** | < 0.0001 | Yes | < 0.0001 | Yes |
| | **5′-pppG vs 5′-OH-G** | 0.54 | No | < 0.0001 | Yes |
| | **5′-OH-A vs 5′-OH-G** | < 0.0001 | Yes | 0.30 | No |
| Synthetic 24 bp dsRNA | **5′-pppA vs 5′-pppG** | < 0.0001 | Yes | 0.70 | No |
| | **5′-pppA vs 5′-OH-A** | < 0.0001 | Yes | < 0.0001 | Yes |
| | **5′-pppG vs 5′-OH-G** | < 0.001 | Yes | < 0.01 | Yes |
| | **5′-OH-A vs 5′-OH-G** | < 0.0001 | Yes | < 0.001 | Yes |

**Supplementary Table S2.** RNA transcripts assessed in the current study.

| Transcript | RNA variant | Sequence, 5′ – 3′ |
|---|---|---|
| **76 bp dsRNA** | 5′-pppA | AGCAAAAGCAGGGUGACAAAGACAUAAUGGAUCCAAACACUGUGUCAAGCUUUCAGGUAGAUUGCUUUCUUUGGC |
| | 5′-pppG | GGCAAAAGCAGGGUGACAAAGACAUAAUGGAUCCAAACACUGUGUCAAGCUUUCAGGUAGAUUGCUUUCUUUGGC |
| **76 bp dsRNA antisense** | antisense to 5′-pppA | AGCCAAAGAAAGCAAUCUACCUGAAAGCUUGACACAGUGUUUGGAUCCAUUAUGUCUUUGUCACCCUGCUUUUGC |
| | antisense to 5′-pppG | GGCCAAAGAAAGCAAUCUACCUGAAAGCUUGACACAGUGUUUGGAUCCAUUAUGUCUUUGUCACCCUGCUUUUGC |
| **24 bp dsRNA** | 5′-pppA | AGCAAAAGCAGGGUGACAAAGACA |
| | 5′-pppG | GGCAAAAGCAGGGUGACAAAGACA |
| **24 bp dsRNA** | antisense to 5′-pppA | UGUCUUUGUCACCCUGCUUUUGCU |
| | antisense to 5′-pppG | UGUCUUUGUCACCCUGCUUUUGCC |
| **pre-let7a** | 5′-pppA | AGUGAGGUAGUAGGUUGUAUAGUUUUAGGGUCACACCCACCACUGGGAGAUAACUAUACAAUCUACUGUCUUUC |
| | 5′-pppG | GGUGAGGUAGUAGGUUGUAUAGUUUUAGGGUCACACCCACCACUGGGAGAUAACUAUACAAUCUACUGUCUUUC |
| **pre-let7a antisense** | antisense to 5′-pppA | GAAAGACAGUAGAUUGUAUAGUUAUCUCCCAGUGGUGGGUGUGACCCUAAAACUAUACAACCUACUACCUCACU |
| | antisense to 5′-pppG | GAAAGACAGUAGAUUGUAUAGUUAUCUCCCAGUGGUGGGUGUGACCCUAAAACUAUACAACCUACUACCUCACC |
| **Y5 WT – tail** | 5′-pppA | AGUUGGUCCGAGUGUUGUGGGUUAUUGUUAAGUUGAUUUAACAUUGUCUCCCCCCACAACCGCGCUUGACUAGCU |
| | 5′-pppG | GGUUGGUCCGAGUGUUGUGGGUUAUUGUUAAGUUGAUUUAACAUUGUCUCCCCCCACAACCGCGCUUGACUAGCC |
| **Y5 WT – tail antisense** | antisense to 5′-pppA | AGCUAGUCAAGCGCGGUUGUGGGGGGAGACAAUGUUAAAUCAACUUAACAAUAACCCACAACACUCGGACCAACU |
| | antisense to 5′-pppG | GGCUAGUCAAGCGCGGUUGUGGGGGGAGACAAUGUUAAAUCAACUUAACAAUAACCCACAACACUCGGACCAACC |
| **Y5 GC clamp – tail** | 5′-pppA | AGCGGGGCCGAGUGUUGUGGGUUAUUGUUAAGUUGAUUUAACAUUGUCUCCCCCCACAACCGCGCUUGCCCCGCU |
| | 5′-pppG | GGCGGGGCCGAGUGUUGUGGGUUAUUGUUAAGUUGAUUUAACAUUGUCUCCCCCCACAACCGCGCUUGCCCCGCC |
| **Y5 GC clamp – tail antisense** | antisense to 5′-pppA | AGCGGGGCAAGCGCGGUUGUGGGGGGAGACAAUGUUAAAUCAACUUAACAAUAACCCACAACACUCGGCCCCGCU |
| | antisense to 5′-pppG | GGCGGGGCAAGCGCGGUUGUGGGGGGAGACAAUGUUAAAUCAACUUAACAAUAACCCACAACACUCGGCCCCGCC |
| **synthetic 11 bp RIG-I ligand** | details provided in the patent WO/2025/088117A1 | |

**Supplementary Table S3.** Primers for IVT of RNA. Yellow highlights T7 promoter, bold shows the modified pair of nucleotides.

| Transcript | RNA variant | Forward primer, 5′-3′ | Reverse primer, 5′-3′ |
|---|---|---|---|
| **short viral** | 5′-pppA | TAATACGACTCACTATT**A**GCAAAAGCAGGGTGACAA | **A**GCCAAAGAAAGCAATCTACCTG |
| | 5′-pppG | TAATACGACTCACTATA**G**GCAAAAGCAGGGTGACAA | **G**GCCAAAGAAAGCAATCTACCTG |
| **short viral antisense** | antisense to 5′-pppA | AAGCTAATACGACTCACTATT**A**GCCAAAGAAAGCAATCTACC | **A**GCAAAAGCAGGGTGACAAAGAC |
| | antisense to 5′-pppG | AAGCTAATACGACTCACTATA**G**GCCAAAGAAAGCAATCTACC | **G**GCAAAAGCAGGGTGACAAAGAC |
| **pre-let7a** | 5′-pppA | TAATACGACTCACTATAAGTGAGGTAGTAGGTTGTATA | GAAAGACAGTAGATTGTATA |
| | 5′-pppG | TAATACGACTCACTATAGGTGAGGTAGTAGGTTGTATA | GAAAGACAGTAGATTGTATA |
| **pre-let7a antisense** | antisense to 5′-pppA | AAGCTAATACGACTCACTATAGAAAGACAGTAGATTGTATAGTTATC | AGTGAGGTAGTAGGTTGTATAG |
| | antisense to 5′-pppG | AAGCTAATACGACTCACTATAGAAAGACAGTAGATTGTATAGTTATC | GGTGAGGTAGTAGGTTGTATAG |
| **Y5 WT – tail** | 5′-pppA | AAGCTAATACGACTCACTATT**A**GTTGGTCCGAGTGTTGTGGGTTAT | **A**GCTAGTCAAGCGCGGTTGT |
| | 5′-pppG | AAGCTAATACGACTCACTATA**G**GTTGGTCCGAGTGTTGTGGGTTAT | **G**GCTAGTCAAGCGCGGTTGTG |
| **Y5 WT – tail antisense** | antisense to 5′-pppA | AAGCTAATACGACTCACTATT**A**GCTAGTCAAGCGCGGTTGTG | **A**GTTGGTCCGAGTGTTGTGG |
| | antisense to 5′-pppG | AAGCTAATACGACTCACTATA**G**GCTAGTCAAGCGCGGTTGTG | **G**GTTGGTCCGAGTGTTGTGG |
| **Y5 GC clamp – tail** | 5′-pppA | AAGCTAATACGACTCACTATT**A**GCGGGGCCGAGTGTTGTGGGTTATTGTT | **A**GCGGGGCAAGCGCGGTTGTGGGGGGA |
| | 5′-pppG | AAGCTAATACGACTCACTATA**G**GCGGGGCCGAGTGTTGTGGGTTATTGTT | **G**GCGGGGCAAGCGCGGTTGTGGGGGGA |
| **Y5 GC clamp – tail antisense** | antisense to 5′-pppA | AAGCTAATACGACTCACTATT**A**GCGGGGCAAGCGCGGTTGTGGGGGGA | **A**GCGGGGCCGAGTGTTGTGGGTTATTGTT |
| | antisense to 5′-pppG | AAGCTAATACGACTCACTATA**G**GCGGGGCAAGCGCGGTTGTGGGGGGA | **G**GCGGGGCCGAGTGTTGTGGGTTATTGTT |

**Supplementary Table S4.** Segmented gradient for LC-MS analysis.

| Time [min] | % A | % B |
|---|---|---|
| **0** | 100 | 0 |
| **1.50** | 100 | 0 |
| **5** | 93 | 7 |
| **11.25** | 85 | 15 |
| **15.75** | 60 | 40 |
| **16** | 0 | 100 |

**Supplementary Table S5.** Primary antibodies.

| Antigen | Vendor | Code | Dilution ratio |
|---|---|---|---|
| **RIG-I** | Cell Signaling Technology | 3743 | 1/1000 |
| **pIRF3** | Cell Signaling Technology | 4947 | 1/1000 |
| **IRF3** | Proteintech | 11312-1-AP | 1/1000 |
| **NUDT16** | Proteintech | 12889-1-AP | 1/1000 |
| **RAN** | Cell Signaling Technology | 4462 | 1/1000 |
| **RANBP1** | Cell Signaling Technology | 8780 | 1/1000 |
| **DHX9** | Proteintech | 17721-1-AP | 1/1000 |
| **α-tubulin** | Proteintech | 11224-1-AP | 1/4000 |
| **PKR** | Proteintech | 18244-1-AP | 1/1000 |

**Supplementary Table S6.** Primers for qRT-PCR.

| Transcript | Forward primer, 5′-3′ | Reverse primer, 5′-3′ |
|---|---|---|
| **Y1** | CTGGTCCGAAGGTAGTGA | TAGTCAAGTGCAGTAGTGAGAAGGG |
| **Y3** | CTGGTCCGAGTGCAGTGGT | TAGTCAAGTGAAGCAGTGGGAGT |
| **Y4** | CCGATGGTAGTGGGTTATCAGAAC | GTCAAATTTAGCAGTGGGGGGTTG |
| **Y5** | TCCGAGTGTTGTGGGTTATTG | TAGTCAAGCGCGGTTGTGG |
| **GAPDH** [1] | AATCCCATCACCATCTTCCA | TGGACTCCACGACGTACTCA |
| **7SL** | GGAGTTCTGGGCTGTAGTGC | TTTGACCTGCTCCGTTTCCG |
| **IPO8** [2] | GGCATACAGTTTAACCTGCCAC | CAGGAGAGGCATCATGTCTGTAA |
| **IFN1β** | ACGCCGCATTGACCATCTAT | TGGCCTTCAGGTAATGCAGA |
| **ISG15** | CGCAGATCACCCAGAAGAT | GCCCTTGTTATTCCTCACCA |
